# Insights into gallbladder cancer pathogenesis from a living organoid gallbladder cholangiocyte biorepository

**DOI:** 10.1101/2025.05.01.651639

**Authors:** Ankita Dutta, Nandita Chowdhury, Akshaya Vijayan Selvarajan, Pritha Banerjee, Abhirupa Kar, Shinjini Chandra, Uma Sunderam, Debdutta Ganguli, Trina Dutta, Dipjit Basak, Smrithi Jayashree Satheeshkumar, Anand Sagar Ragate, Shekhar Krishnan, Saugata Sen, Manas Kumar Roy, Sudeep Banerjee, Rajgopal Srinivasan, Paromita Roy, Vaskar Saha, Anindita Dutta, Dwijit GuhaSarkar

## Abstract

Gallbladder cancer (GBC) while rare worldwide has a high prevalence in India. Pathogenesis is unclear and outcomes poor. Gallbladder cholangiocyte organoids (GCOs) or gallbladder carcinoma organoids (GBCOs) were developed and serially propagated from surgically resected gallbladder tissues with benign or malignant diseases, respectively. Patient derived organoids (PDOs) were derived from 15 normal; 58 inflamed; 12 xanthogranulomatous cholecystitis (XGC); 5 pre-invasive neoplasm and 13 invasive malignant gallbladder pathologies. Protocol optimisation achieved 58% (69/119) success in organoid generation and expansion. Organoids maintained tight junction integrity; P-gp pump and enzymatic activity; preserved tissue-specific gene and protein marker expression; histological features and genetic variations. Cryopreserved organoids from 62 patients with primary tissue and high-quality DNA, RNA and protein derivatives have been banked. In gene expression analyses of tissue, XGC samples clustered with malignant subtypes, separate from benign pathologies. Derived XGC organoids showed a similar clustering. Enriched hallmark pathways in XGC support neoplastic change through chronic inflammation. PDOs generated from different gallbladder pathologies are a promising model to investigate the pathogenesis of GBC.

**Graphical Abstract:** 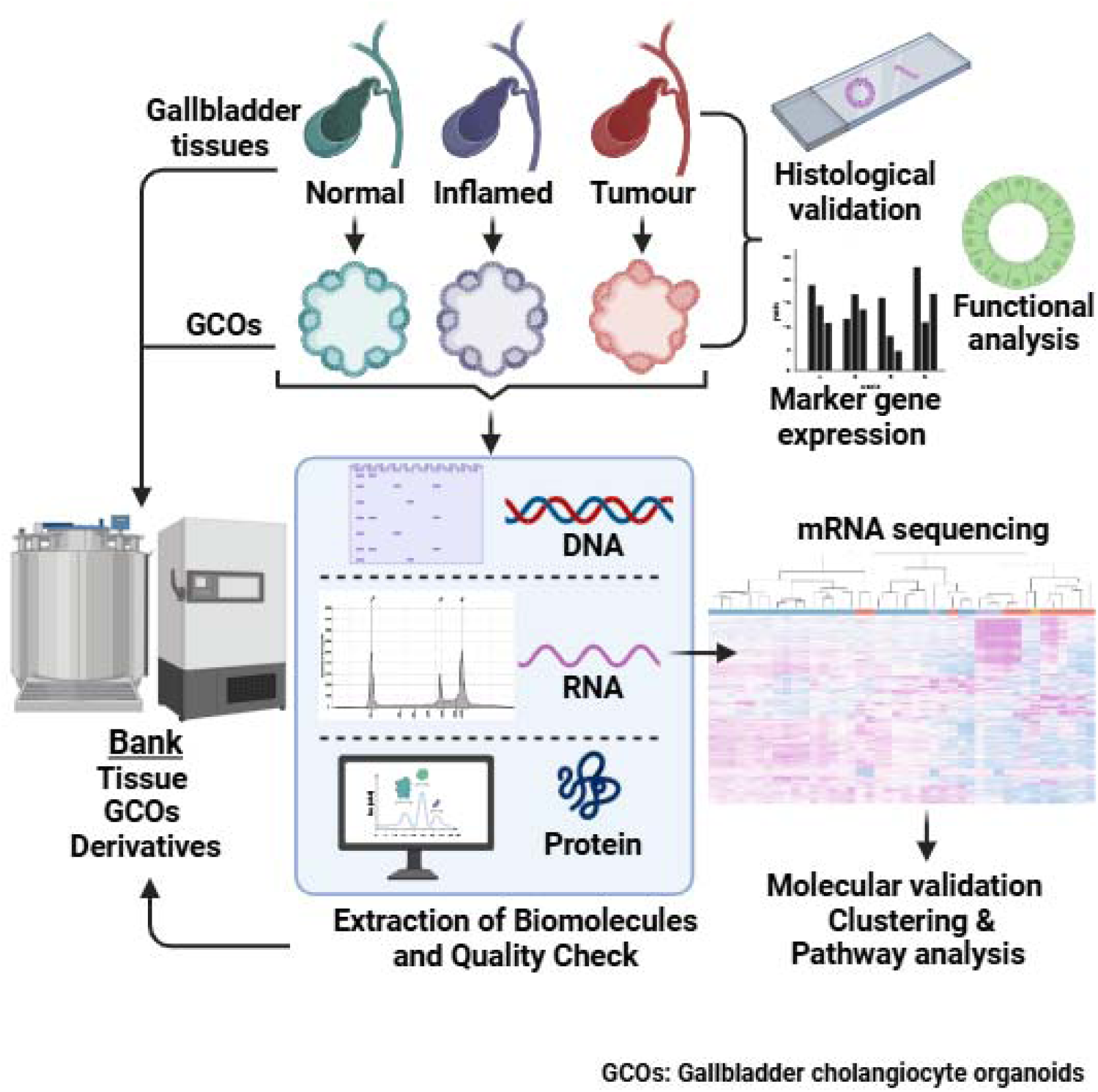

## Introduction

Gallbladder cancer (GBC) is the most common and aggressive form of cancer among biliary tract cancers^1^. It is a neglected cancer as it is rare in high income countries, but has a high incidence in north and north-east India^2^. The aetiology of GBC pathogenesis is not well understood. Gallstones, pro-inflammatory lithogenic bile, bacterial infections, heavy metal toxicity, high blood cholesterol, obesity and female gender have been implicated.^3^ The current hypothesis is that a chronic inflammatory process leads to histogenic progression through either metaplasia-dysplasia-carcinoma or adenoma/pre-invasive neoplasm to an invasive carcinoma.^4^ Most (>90%) patients are diagnosed at a late stage (III/IV) and the 2-year overall survival has remained static at 30% for over 3-decades.^5^

A suitable pre-clinical model is required to better understand pathogenesis and identify curative options. While erbB2 transgenic mice spontaneously develop GBC, the anatomical differences and lack of heterogeneity does not make this a suitable model.^6,7^ Patient derived organoids (PDOs) have been generated from normal and cancerous gallbladders. These organoids retain morphological, histological and functional characteristics of the primary tissue and have been used for drug screening ^8–12^ with moderate success. A possible reason for this is the low success rates in generating gallbladder carcinoma organoids (GBCOs) ^8^ and maintaining these in long-term cultures.^9,11^ When exposed to *Salmonella* sp, human gallbladder cholangiocyte organoids (GCOs) proliferate in spite of double-strand DNA damage ^13^ and murine derived GCOs, on a background of *Tp53* mutation and *c-Myc* amplification, develop cellular features associated with GBC.^14^ This suggests that GCOs/GBCOs generated from different gallbladder pathologies could be suitable models to investigate pathogenesis and generate a pre-clinical model for drug discovery. Here we describe processes by which we have successfully generated and passaged PDOs from normal and a variety of diseased gallbladders including GBC to create a living organoid bank as a resource for the community.

## Results

### Establishing workflows for clinical sample collection, derivative creation and biobanking

To maintain sterility, cellular viability and intracellular molecular integrity, all tissues were collected directly and transported to the laboratory in freshly prepared tissue transport media on ice. Tissue transport medium composition was optimised over time to PBS with epidermal growth factor and antibiotics, reducing cost without compromising cell viability, culture sterility and RNA quality. [Supplemental table S1].

Surgically resected fresh gallbladder tissue, core needle biopsy and formaldehyde fixed paraffin embedded (FFPE) blocks were collected from a total of 357, 17 and 22 patients, respectively. Fresh tissues were prioritised for direct processing to obtain primary cells (n=300) for cryopreservation or organoid derivation. Where available, fresh tissues were stored in RNAlater^TM^ (ThermoFisher Scientific) (n=143); snap frozen (SF) (n=137) and stored at -80°C for DNA, RNA and protein extraction or placed in tissue lysis buffer (ATL buffer, Qiagen) and stored at room temperature for genomic DNA (gDNA) extraction. [Figure 1 and Supplemental Table S2].

**Figure 1.**
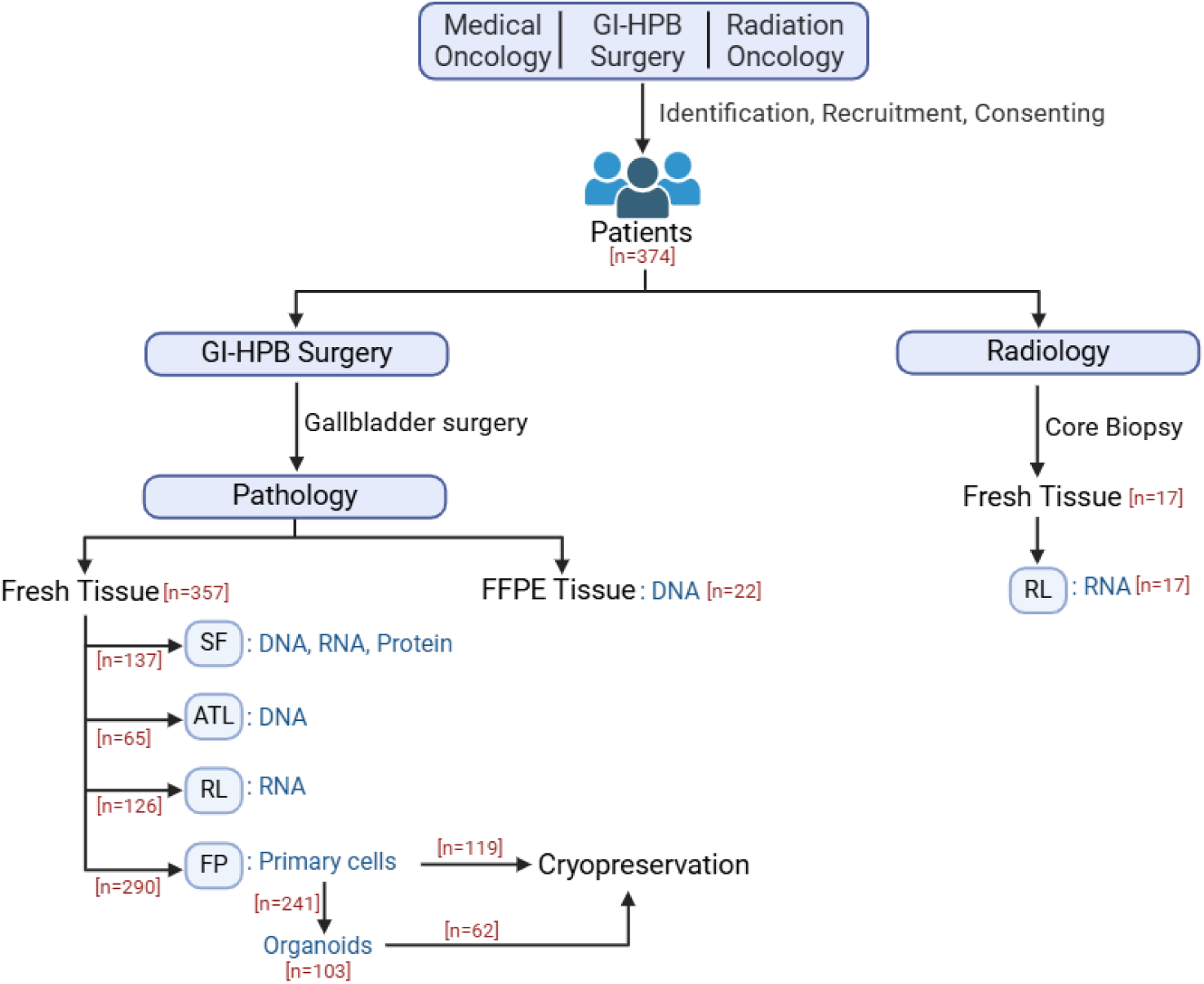
Workflow for patient tracking and biobanking for gallbladder tissues at Tata Medical Centre (TMC). **(A)** Gallbladder (GB) cancer patients presented primarily to the GI-HPB surgery, medical oncology and radiation oncology departments. Diagnosis was confirmed by radiology and pathology. For patients with resectable GB cancer or non-malignant GB disease, or normal GB requiring removal for other reasons, surgical specimens were obtained. For non-resectable disease, image guided core biopsy samples were obtained. Surgical or core biopsy procedure derived fresh GB tissue samples were collected in specified tissue transport media. Fresh tissues were either snap frozen (SF) in liquid nitrogen and stored at - 80°C, or stored in Qiagen tissue lysis buffer (ATL) at room temperature for DNA extraction or in RNAlater^TM^ (RL) at 4°C for RNA extraction. SF tissues were used as a source for either DNA, RNA or protein extraction. Tissues were freshly processed to isolate primary cells and were either directly grown into organoids or cryopreserved for future use. Part of the organoid cultures were cryopreserved and banked. FFPE blocks were used as alternative source for DNA extraction. GI-HPB, Gastro-Intestinal Hepatopancreaticobiliary; FFPE, Formalin-fixed paraffin embedded tissue;

gDNA was successfully obtained from fresh tissue (n=65 stored in ATL and n=8 as SF) and FFPE blocks (n=22); total RNA from surgical tissue (n=126) and core biopsies (n=17) and protein from SF tissues (n=20). Agarose gel electrophoresis confirmed integrity of gDNA extracted from >85% of tissues stored in ATL buffer^TM^ and from >70% of the tissues stored in SF condition or as FFPE blocks [Figure 2A-B]. A minimum of 5 (10 µm each) sections were required to obtain a median yield of >2µg of gDNA from FFPE blocks [Figure 2C-D]. For ATL-stored and SF tissue samples, 24-30mg of starting tissue material yielded a median of >3µg and >4µg of total extracted gDNA per sample, respectively [Figure 2D]. RNA integrity was optimal when tissue samples were collected in PBS and stored in RNAlater^TM^ and extracted using silica column-based RNA-extraction but not when samples were stored or extracted in TRI reagent^TM^ [Figure 2E]. High quality RNA was obtained from both surgical tissue and core biopsies but not from SF tissue [Figure 2F-G]. This could be attributed to the delay between surgical resection and sample freezing timepoints. Liquid chromatography followed by tandem mass spectrometry (LC-MS/MS) showed satisfactory quality of proteins extracted from SF tissues from both malignant and non-malignant pathologies. LC-MS/MS identified a median of 4923 unique proteins from tissues in data dependent acquisition (DDA) mode [Figure 2H-I].

**Figure 2.**
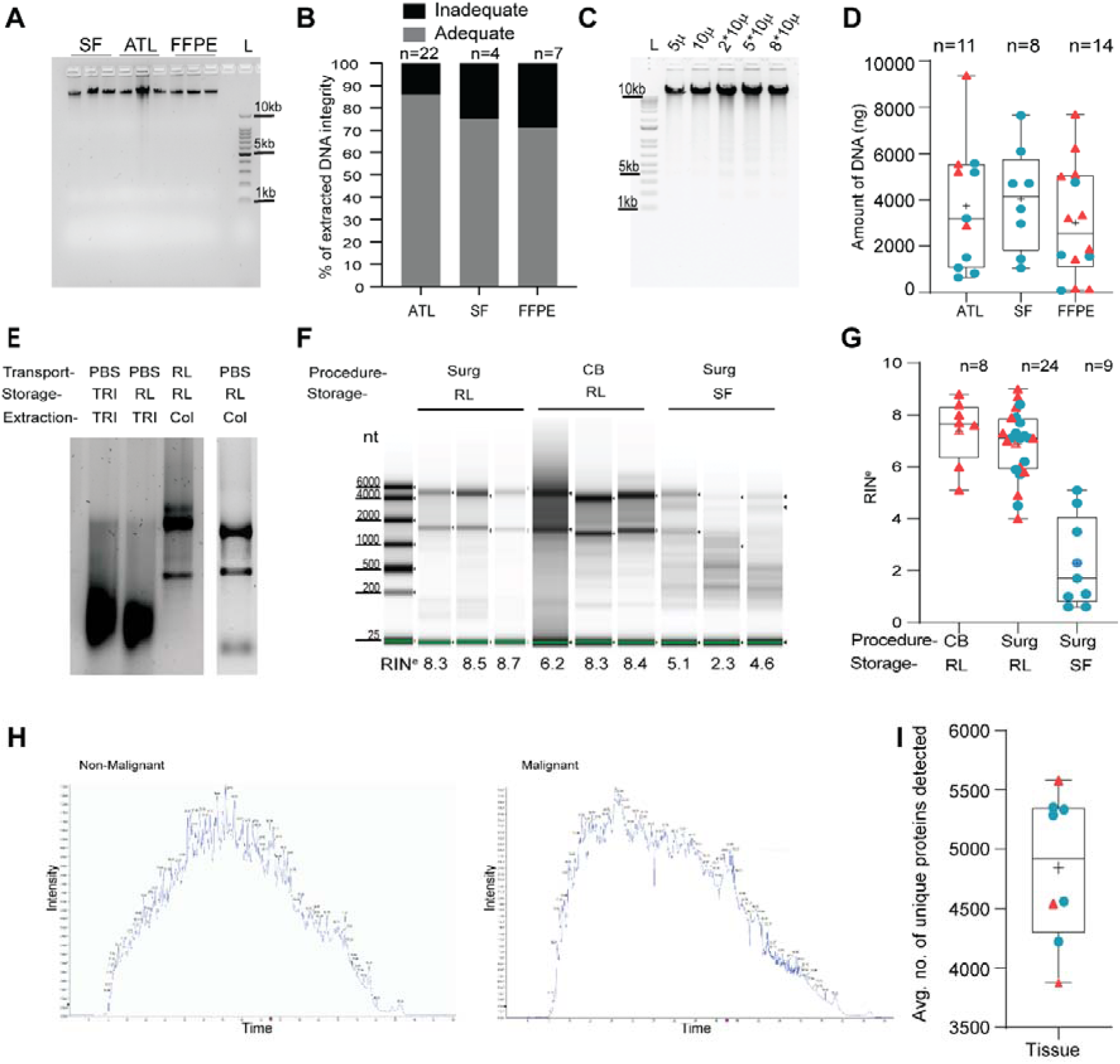
Quality, purity and quantity of the extracted biomolecules were adequate for downstream high throughput experiments. **(A)** Representative image of genomic DNA (gDNA) integrity analysis by 1% agarose gel electrophoresis. Corresponding tissue storage conditions have been indicated above respective lanes. (**B)** Bar graph depicting proportions of tissue gDNA sample integrity according to storage conditions. **(C)** gDNA from FFPE tissue sections of varying thickness and numbers as indicated by respective lanes. **(D)** Box and whisker plots depicting the distribution of total quantity of gDNA extracted from tissues stored in different conditions. DNA extraction was performed from 27± 3 mg of each of FP or SF tissue and 5 pieces of 10µm sections each were used for every individual FFPE sample. **(E)** Representative images of 1% agarose gel electrophoresis analysis of tissue RNA samples showing varying integrity depending on extraction method. Tissue transport media, storage and extraction methods for different samples are indicated above each respective lane. **(F)** Representative images of column (Qiagen RNeasy) purified tissue RNA samples run in automated gel electrophoresis (Agilent 4200 TapeStation System). Tissue collection procedure and storage conditions are indicated above respective lanes. **(G)** Box and whisker plots depicting the distribution of the RIN^e^ values of organoid RNA samples determined in RNA quality analyses by 4200 TapeStation System (Agilent). **(H)** Representative images of intensity-time chromatogram plots from LC-MS/MS analyses (DDA mode) of GB tissues from XGC (left) and AdSqCa pathology (right). **(I)** Box and whisker plot showing the distribution of number of unique proteins detected from each tissue protein sample. Equal amount (100ng) and equal volume (2µl) of eluted gNA were loaded in each lane for agarose gel electrophoresis analyses shown in Figure 2A and 2C, respectively. Equal volume (3µl) of RNA was loaded in each lane for 2E. For the box and whisker plots in **2D**, **G** and **I**, boxes, whiskers, horizontal lines and ‘+’ symbols indicate the interquartile range, minimum to maximum range, median and mean values, respectively. Red triangular and blue circular data points indicate malignant/ preinvasive neoplasm and non-malignant samples, respectively. SF, snap frozen tissue; ATL, Qiagen tissue lysis buffer for DNA extraction; FFPE, formalin fixed paraffin embedded tissue; L, ladder; CB, Core Biopsy; Surg, Surgery; NM, Non-malignant; M, Malignant; PBS, Phosphate Buffered Saline; TRI, TRI reagent^TM^; RL, RNAlater; Col, Column based purification using RNeasy^TM^ kit (Qiagen), RIN^e^, RNA integrity number equivalent. LC-MS/MS, Liquid chromatography followed by tandem mass spectrometry. DDA, Data dependent acquisition. XGC, xanthogranulomatous cholecystitis; AdSqCa, adenosquamous carcinoma.

### Protocol optimisation for the development and expansion of patient derived GCO lines from malignant and non-malignant gallbladder tissues

PDO lines were successfully grown from both malignant and non-malignant gallbladder mucosal cholangiocytes. Isolation of cholangiocytes from fresh surgical tissues were done by mechanical scraping when sufficient tissue was available. When tissue amount was limited, enzymatic (collagenase/dispase) digestion was used to release cholangiocytes [Figure 3A]. PDOs were successfully generated from different gallbladder tissue pathologies using both the cell isolation methods [Figure 3B]. Two different growth media were used, adapted from previous reports, type A ^9^ and type B ^15^ [Supplemental Table S3]. Organoids were derived and propagated in both types of culture media [Figure 3C]. Both GCO and GBCO cultures were expanded over several (>25) passages [Figure 3D]. We observed that GCOs had regular cystic structures with translucent central lumens. In contrast, GBCOs often had irregular and optically dense structures or a mixed phenotype [Figure 3C-D]. Enzymatic digestion (TrypLE Express^TM^) was superior to mechanical passaging [Figure 3E], and enabled long term maintenance (≥ 10 passages) of organoid cultures [Figure 3F].

**Figure 3.**
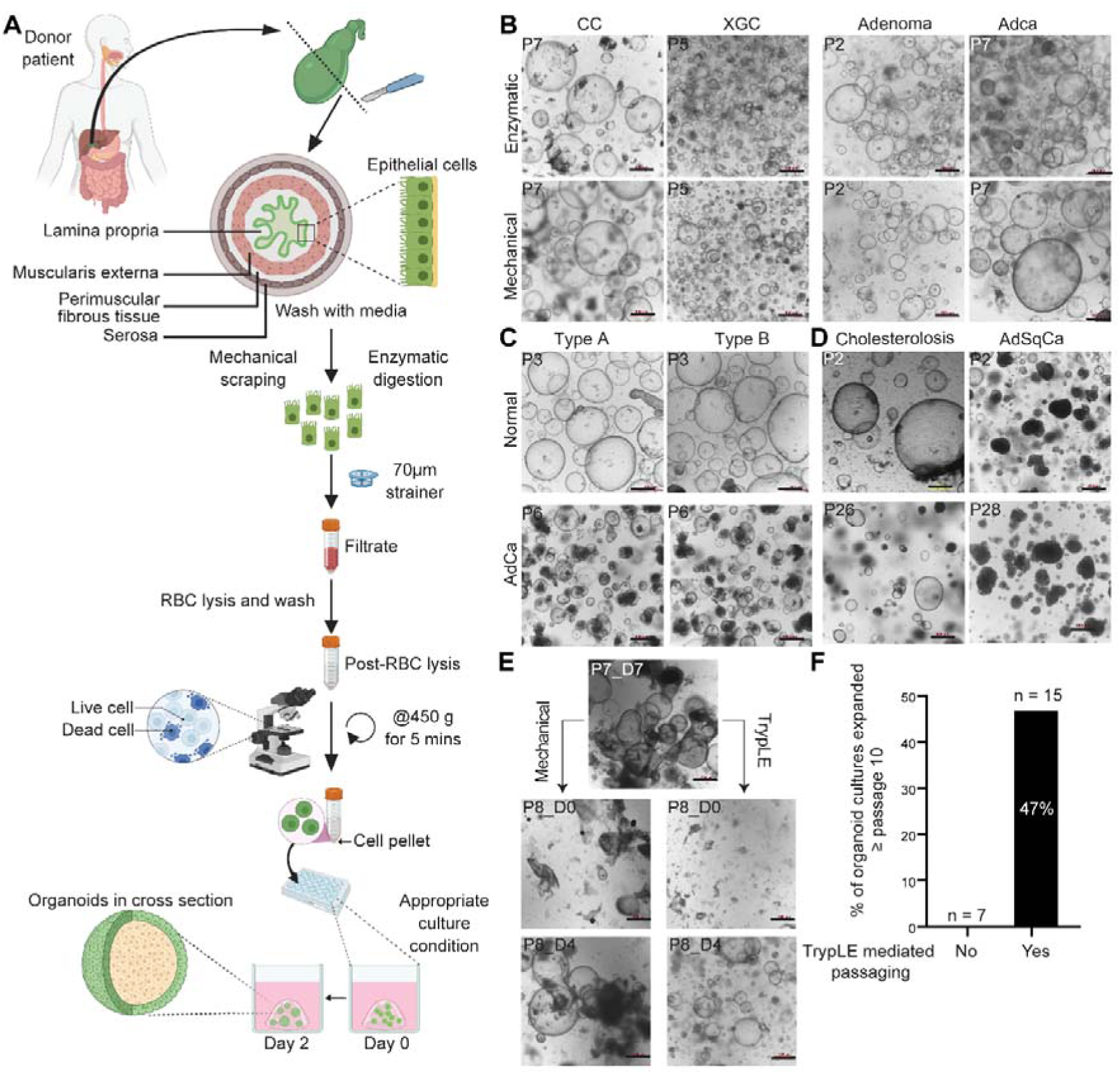
Organoids developed from patient tissues could be maintained long-term in culture. **(A)**Schematic workflow of gallbladder tissue processing for developing organoids. **(B)** Representative images of organoids developed from non-malignant inflamed (CC and XGC), pre-invasive neoplastic (adenoma) and malignant (AdCa) gallbladder epithelial cells isolated by enzymatic dissociation (top panel) and mechanical scraping (bottom panel). **(C)** Representative images of GCOs and GBCOs developed from normal and AdCa gallbladder pathology in Type A (left) and Type B (right) culture media. **(D)** Representative images of organoids grown from non-malignant (cholesterolosis) and malignant (AdSqCa) tissues maintained over 26 and 28 passages, respectively. **(E)** Representative images of a degenerating GCO (CC) culture (top). Passaging by TrypLE enzymatic digestion (below right) showed better growth than when passaged by mechanical shearing (below left). Passage numbers and days after last passages are indicated in the respective images. **(F)** Bar plot depicting the percentage of organoid cultures reaching passage number ≥ 10 before or after introduction of TrypLE digestion mediated passaging for degenerating cultures. Scale bar: 500μm. P, passage number. D, days since last passage. CC, chronic cholecystitis. XGC, xanthogranulomatous cholecystitis; AdCa, adenocarcinoma; AdSqCa, adenosquamous carcinoma; GCO, gallbladder cholangiocyte organoids; GBCO, gallbladder carcinoma organoids. Microscope: Nikon TS2-FL, bright field mode.

Primary cells were isolated from the mucosal layer of freshly collected gallbladder tissues from 290 participating patients undergoing cholecystectomy between September, 2019 and December, 2023. Organoid cultures were initiated from 241 patients (surgical sample), of which PDOs could be developed and propagated successfully (>passage 4) from 103 samples. While mean success in organoid generation and expansion (> passage 4) from surgically resected tissues over the full period of the study was 42.7%, success improved from 20% (n= 59) in the period of 2019-20 to 58% (n=119) in 2022-23 with continuous optimisation. Attempts to grow organoids from needle core biopsy tissues were unsuccessful (n=10) [Supplemental Table S2].

### Protocol optimisation for biomolecule extraction from GCOs

Integrity of gDNA samples extracted from GCOs/GBCOs was confirmed by agarose gel electrophoresis [Supplemental Figure S1A-B]. Organoids from two confluent wells of a 24-well culture dish yielded a median of 2.7µg of gDNA [Supplemental Figure S1C]. Unlike primary tissues, total RNA extracted from PDOs both by silica column (RNeasy Micro^TM^ kit) and TRI-reagent extraction methods maintained adequate integrity with median RNA integrity (RIN^e^) values of 8.8 and 9.4, respectively [Supplemental Figure S1 D-F]. Extracted proteins both from GCOs and GBCOs could be used successfully for LC-MS/MS analysis, resulting in detection of a median of 5107 unique proteins in DDA mode [Supplemental Figure S1G-H].

### Organoids could be developed and retrieved from cryopreserved primary cells and previously grown organoids, respectively

Gallbladder PDOs were successfully grown and expanded over several passages from cryopreserved primary cholangiocytes [Figure 4A]. Organoids were further cryopreserved either as whole organoids (average organoid diameter <200µm) or after digestion with TrypLE Express^TM^. By both methods, cryopreserved organoids were successfully retrieved and expanded over multiple passages retaining structures observed at primary culture [Figure 4B]. Both primary and cryo-retrieved organoids preserved histological features of the source tissues [Figure 4C]. We found that DNase I treatment immediately upon thawing was essential for the development of organoids from cryopreserved primary cells [Figure 4D]. The addition of the Rho-associated kinase inhibitor (Y-27632 dihydrochloride) in the thawing media and post-retrieval culture media (for the first 48 hrs) improved the retrieval success with an odds ratio of 33 [z-score = 2.82; p = 0.002].

**Figure 4.**
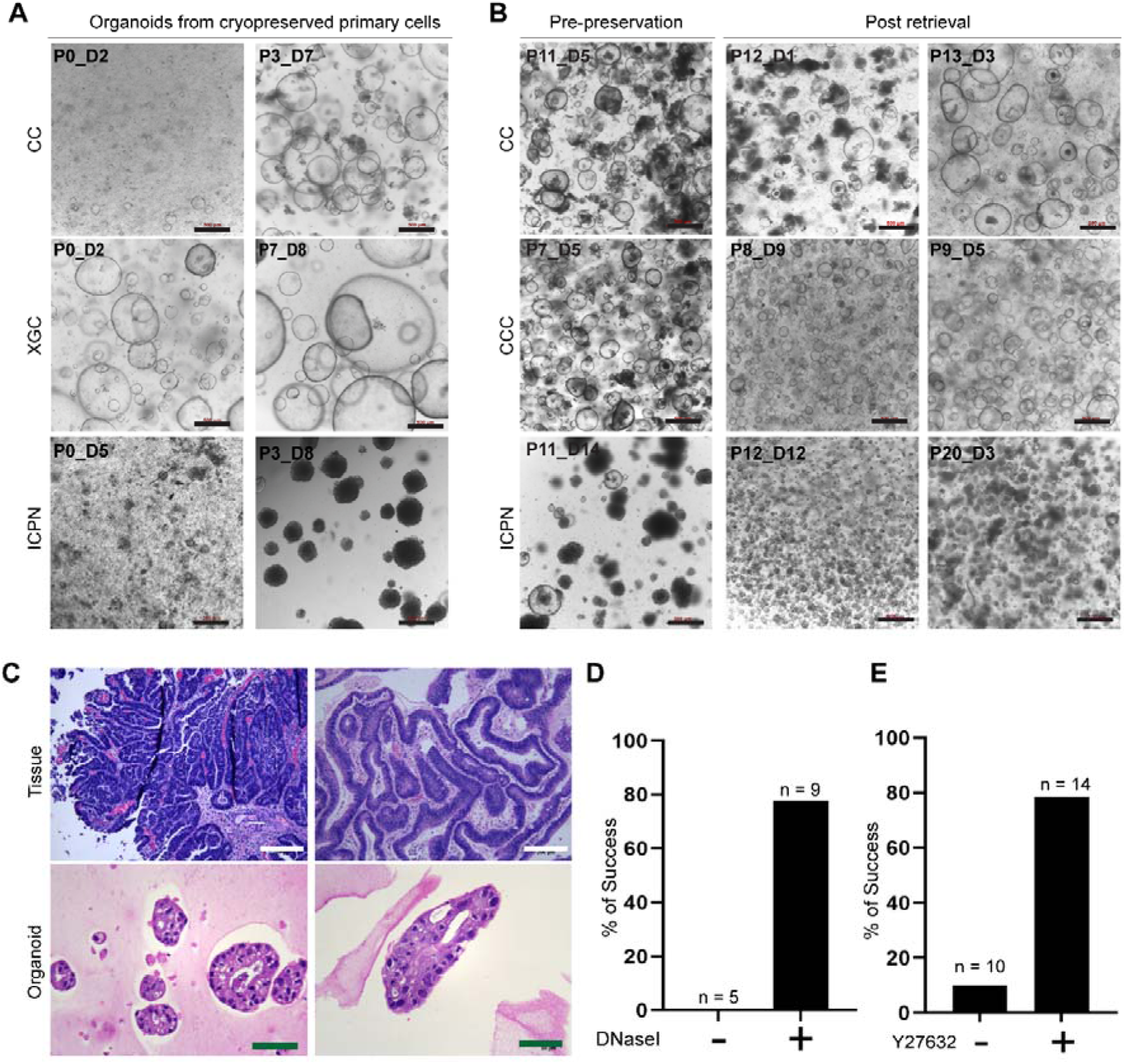
Organoids could be developed successfully from cryopreserved primary cells and organoids. **(A)** Representative images of developed organoids from cryopreserved primary cells maintained over several passages. Source tissue pathologies are indicated on the left. **(B)** Representative images of organoids regrown upon retrieval of cryo-preserved whole organoids (top row) or TrypLE digested organoids (bottom two rows). Pathologies of the source tissues are indicated on the left. **(C)** Representative images of HE-stained FFPE tissue sections from ICPN pathology (top) showing dysplastic cytological features were preserved in organoids (bottom) from the corresponding source tissues. These organoids were grown from cryopreserved primary cells (left bottom) or cryo-preserved organoids (right bottom). **(D)** Bar plots showing proportion of successful development (green) of organoid cultures from cryopreserved primary cells (left) or banked GCOs (right) in presence or absence of DNase-I treatment or use of ROCK inhibitor (Y-27632 dihydrochloride) post-retrieval, respectively. Scale bar: 500μm (red), 200μm (white), 50μm (green). P, passage number of cultures; D, days after last passage; CC, Chronic cholecystitis; CCC, Chronic cholecystitis with cholesterolosis; ICPN, Intracholecystic papillary-tubular neoplasm. HE. Haematoxylin and Eosin. Microscope: Leica DMi8 (HE sections) and Nikon TS2-FL (whole organoids), bright field mode.

One hundred seven living gallbladder PDO lines, from 62 patients have been banked. Gallbladder pathologies included invasive malignant (n= 9), pre-invasive neoplasms (n=5), XGC (n=11) and normal or other non-malignant diseases (n=37) [Supplemental table S2].

### GCO and GBCO lines preserved histological, functional and molecular features of the source tissue

Gallbladder PDO lines were analysed histologically to evaluate preservation of architectural and cytopathological features of the original tissue samples. Given the small size and fragility, formalin-fixed organoids were trapped in eosin labelled coagulum prior to the generation of FFPE blocks [Figure 5A and Methods]. Haematoxylin-Eosin (HE) stained FFPE sections showed that GCO lines had systematically organised single layer of epithelial cells surrounding a central lumen along with normal nuclear morphology reflecting their parent tissues. Mucin secreting glands were observed in some samples. Compared to other non-malignant samples, GCO lines derived from XGC tissues showed higher nucleus to cytoplasm ratio (N/C) and cribriform structures. GBCO lines developed from ICPN as well as AdCa showed cytological and architectural features indicative of moderate to high grade of dysplasia. Multi-layering of cells forming cribriform structures or filling the central lumen were observed. Cells were hyperchromatic with high N/C and frequently with prominent multiple nucleoli, clumped chromatin, nuclear disarray and atypia [Supplemental Figure S2 and S3]. Overall, the histological analysis confirmed that gallbladder PDO lines preserved architectural and cytopathological features of the corresponding patient tissues [Figures 5B].

**Figure 5:**
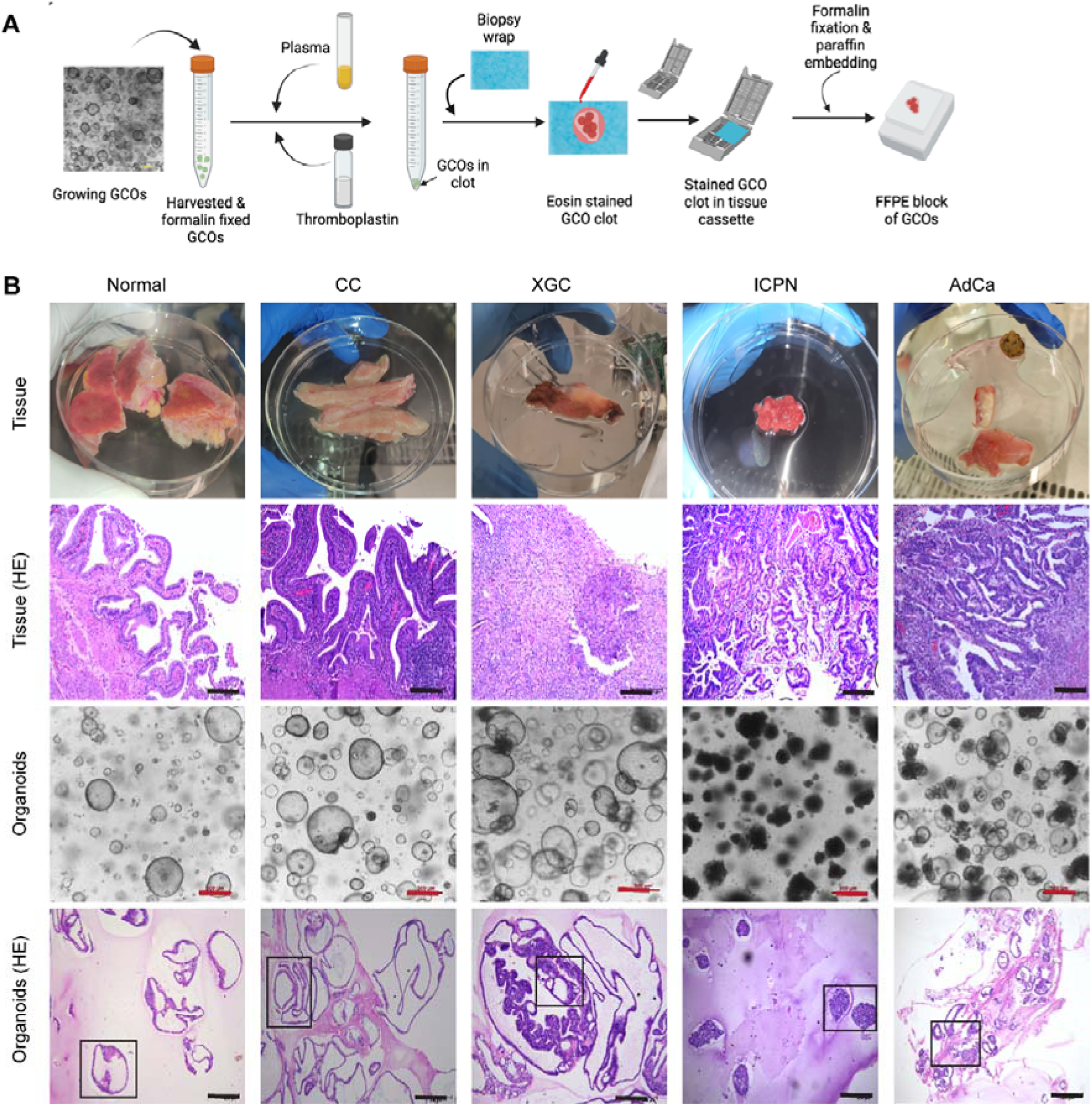
Organoids developed from different gallbladder tissue pathologies preserved the histological features of the parent tissues. **(A)** Schematic workflow for the preparation of organoid FFPE blocks. **(B)** Images showing the fresh tissue samples received at the laboratory and their respective HE stained FFPE sections (top two horizontal panels, respectively). The corresponding pathology is indicated above each vertical panel. Bright field microscopic images showing the corresponding organoids and their respective HE stained FFPE sections (bottom two horizontal panels, respectively). Scale bars: 200µm (black), 500µm (red). Higher magnification images of the areas marked in the bottom row have been shown in Supplemental Figure S2. FFPE, formalin fixed paraffin embedded; CC, chronic cholecystitis; XGC, xanthogranulomatous cholecystitis; ICPN, intracholecystic papillary-tubular neoplasm; AdCa, adenocarcinoma; HE, haematoxylin-eosin. Microscope: Leica DMi8, 10X air objective (HE sections) and Nikon TS2-FL, 4X air objective (whole organoids), bright field mode.

Organoids showed activity of alkaline phosphatase [Figure 6A] and gamma glutamyl transferase [Figure 6B], expressed *KRT7*, *KRT19, CFTR* and *SOX9*, but not *AFP* or *ALB* which are primarily expressed in hepatocytes. [Figure 6C]. Immunofluorescence staining of organoids confirmed expression of MUC5B and CK19 proteins [Supplemental Figure S4A] consistent with previous reports^16,17^. Intense diffuse expression of CK7 in all tissues and corresponding GCO/GBCO lines were observed with IHC. CK20 expression was variable in intensity and distribution across tissues, concordant in 6/8 corresponding PDO lines [Figure 6D, E and Supplemental Figure S4B]. HER2 overexpression has been reported in 12-17% of GBC patients^18^. Out of the 9 patients tested for HER2, expression was positive (3+) and equivocal (2+) for 1 and 3 patient tissues respectively, rest had no expression. But expression was not detected in any of the corresponding GBCO lines. Morphologically the GBCOs screened retained the malignant phenotype. Possibly the lack of detectable HER2 reflects spatial heterogeneity in tissue expression [Supplemental Figure S4C] and/or the different microenvironment in culture. Functionally, on exposure to FITC labelled dextran (4kDa), there was no luminal uptake of the molecules in the GCOs. Fluorescent signal from FITC was visible in the lumen upon exposure to the Ca++ ion chelating agent ethylene glycol-bis (β-aminoethyl ether)-N,N,N′,N′-tetraacetic acid (EGTA). This confirmed the presence of active tight junctions forming an intact epithelial barrier in the GCOs [Figure 6F]. The fluorescent dye Rhodamin123, which is a substrate of the ATP-binding cassette transporter P-glycoprotein (P-gp) pump (encoded by *ABCB1*), enters the lumen of GCOs when added in culture. This active uptake was inhibited by P-gp inhibitor verapamil hydrochloride [Figure 6G], confirming that GCO cholangiocytes preserve active P-gp mediated transport across the epithelial barrier.^19,20^ Only GCOs (derived from CC and cholesterolosis) were subjected to the tight junction and P-gp pump activity assay as GBCO lines had a partial or complete absence of lumen and thus were not suitable for luminal uptake assays. Taken together, we conclude that the developed gallbladder PDOs represented the parent tissues histologically, functionally and at the molecular marker expression level.

**Figure 6:**
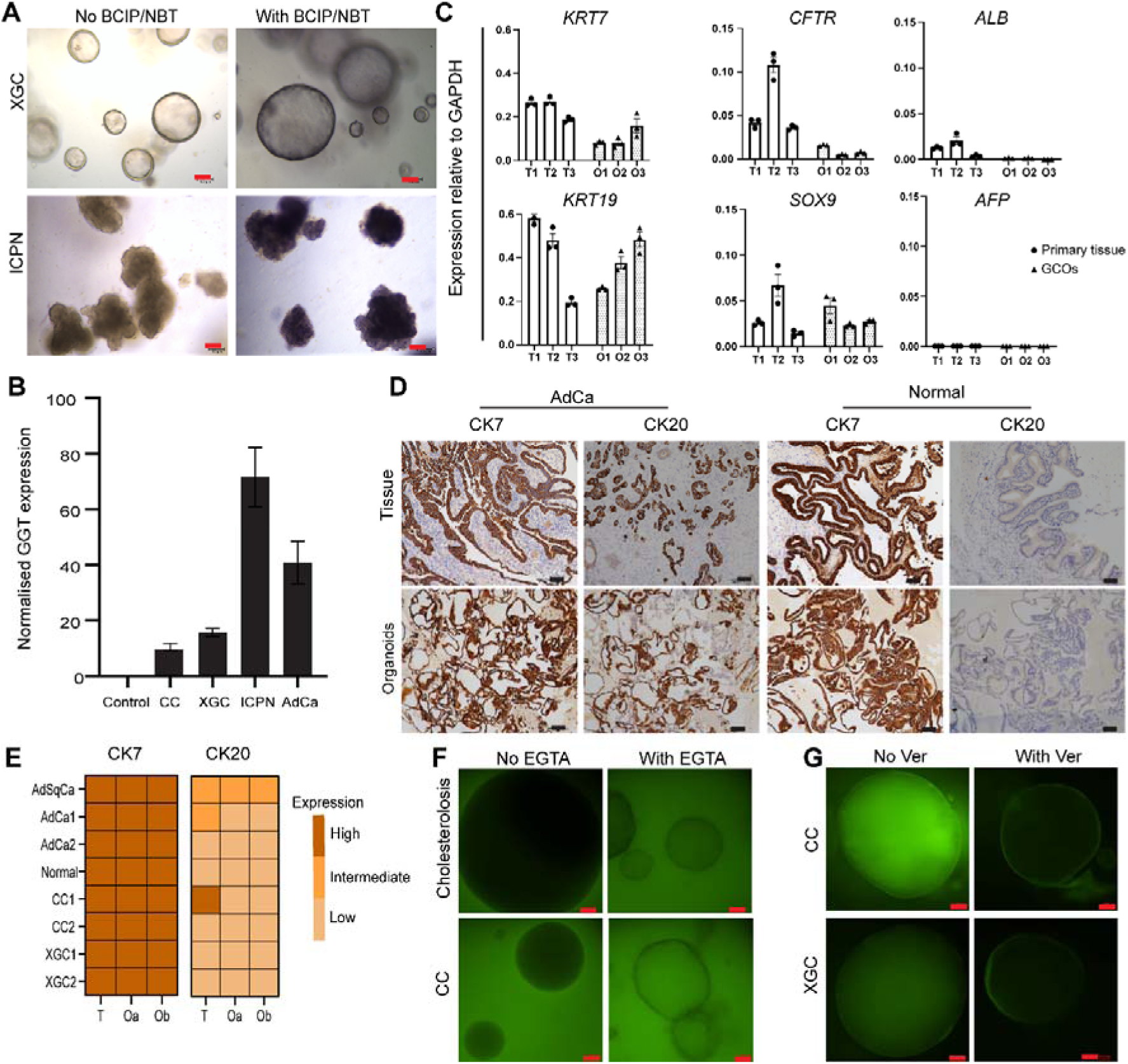
Organoid lines developed from different tissue pathologies retained functional activity and preserved expressions of the molecular markers at RNA and protein level. **(A)** Representative images showing activity of ALP enzyme in organoids derived from ICPN and XGC tissues. **(B)** GGT activity was detected from organoids derived from GB tissues with different pathologies. **(C)** q-RT-PCR confirms the cholangiocyte lineage through the expression of *KRT7, KRT19, CFTR*, *SOX9, ALB* and *AFP* in primary tissues and their corresponding GCOs (n = 3). Error bars show Mean +/-SEM. O1, O2 and O3 indicate corresponding GCO lines grown from tissues T1, T2 and T3 derived from chronic cholecystitis with cholestrolosis patients n196, n198 and n33, respectively. **(D)** Representative images of the gallbladder tissues and corresponding organoid lines showing similar level of expression of the marker proteins CK7 and CK20. **(E)** Heatmap showing comparative expression of KRT7 and KRT20 markers in tissues and two corresponding organoid lines for each patient as assessed from IHC analysis. See methods for scoring details. Oa and Ob indicate organoid lines grown in type A and type B media from the tissue, respectively. **(F)** Representative fluorescence microscopy images of FITC-Dextran permeability assay displaying the retention of functional tight junction and **(G)** rhodamine 123 active transport assay showing P-gp pump activity. Scale bars: 100µm. Microscope: Leica DMi8 (A, D) and Nikon TS2-FL (F, G). GCOs, gallbladder cholangiocyte organoids; CC, chronic cholecystitis; AdCa, adenocarcinoma; AdSqCa, adenosquamous carcinoma; XGC, xanthogranulomatous cholecystitis; ICPN, intracholecystic papillary-tubular neoplasm; EGTA, ethylene glycol-bis (β-aminoethyl ether)-N, N, N′, N′-tetraacetic acid; Ver, Verapamil hydrochloride; BCIP, 5-bromo-4-chloro-3-indolyl phosphate; NBT, nitro-blue tetrazolium; ALP, Alkaline phosphatase; GGT, Gamma-glutamyltransferase; FITC, Fluorescein isothiocyanate; T, tissue; O, organoid;

### Transcriptomic analysis demonstrates suitability of the developed gallbladder organoids for studying disease biology

Transcriptomic analyses of paired primary GB samples and derived PDOs showed 94-96% of the single nucleotide polymorphisms (SNP) were retained by the PDOs. SNPs were preserved in multiple organoid lines developed from a single primary tissue irrespective of different tissue dissociation methods, culture conditions or passages [Figure 7A].

**Figure 7:**
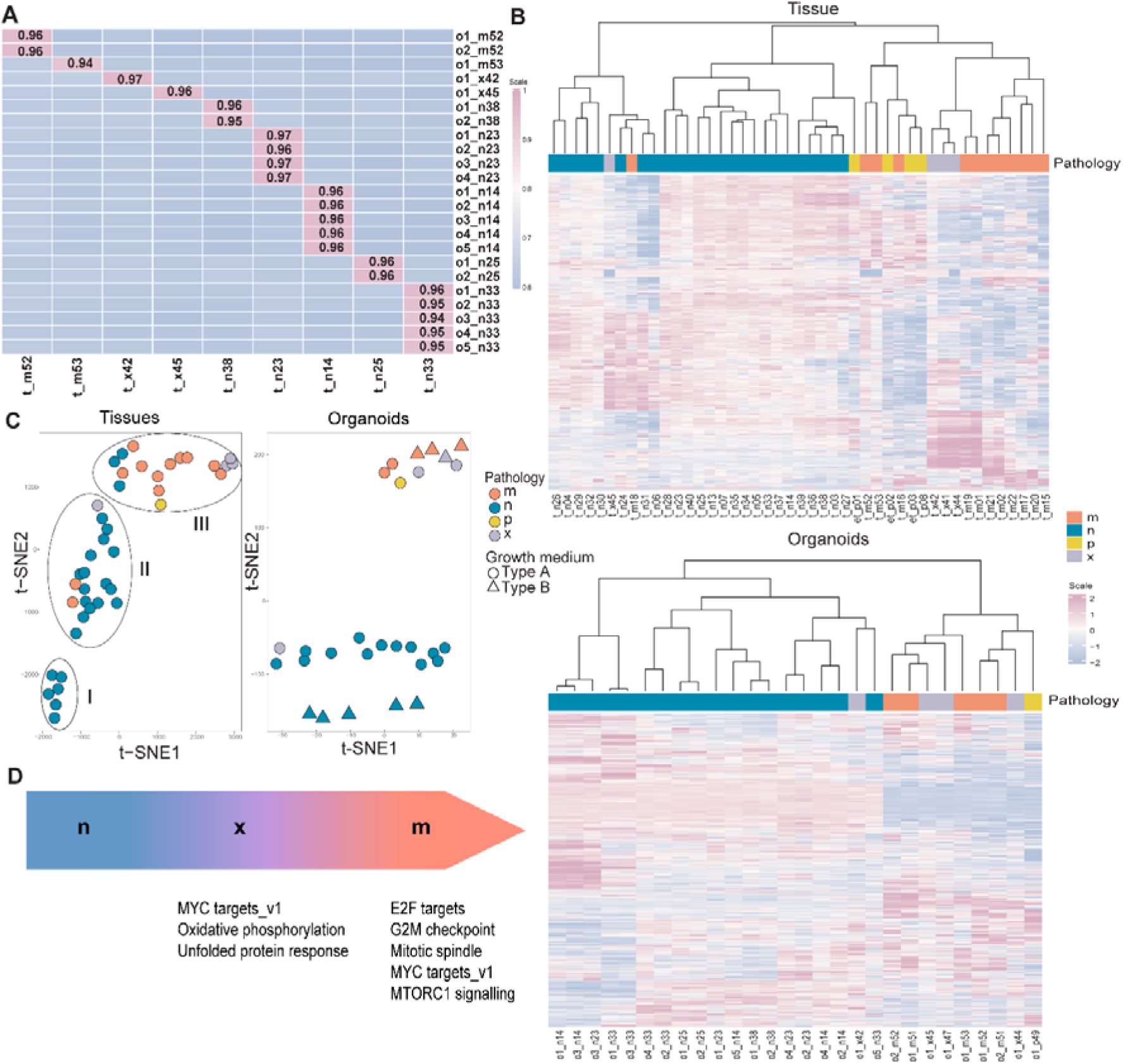
Transcriptome analysis demonstrates suitability of developed organoids for studying gallbladder pathogenesis. **(A)** Correlation matrix showing the fraction of the shared single nucleotide polymorphism (SNP) sequences between the organoid lines (Y-axis) and the patient tissues (X-axis). Variants were called from the two-pass aligned RNA-Seq bam with the haplotyper caller from GATK. **(B)** Unsupervised hierarchical clustering based on the scaled vst counts of the gene expression profile of tissues (top) and organoids (bottom). Hierarchical clustering (with Pearson correlation and linkage clustering algorithm) was performed on top 1000 highly variable genes. For gene names, corresponding raw counts and vst transformed data of the 1000 highly variable genes of tissues and organoids see the Supplemental Tables S4A, S4B, S5A and S5B. **(C)** t-SNE clustering based on the gene expression profile of tissues (left panel) and organoids (right panel). t-SNE clustering was performed on top 1000 highly variable genes. Pathology groups have been indicated by the colours and organoid growth media types by the shapes. **(D)** Proposed model of GBC pathogenesis with XGC as an intermediate state between normal and malignant GB. Enriched Hallmark pathways in x and m group pathologies have been indicated respectively. n, normal or inflamed gallbladder pathology except XGC; x, xanthogranulomatous cholecystitis (XGC); p, pre-invasive neoplasms (Intracholecystic papillary-tubular neoplasm or adenoma); m, invasive malignant pathology (adenocarcinoma or adenosquamous carcinoma); t, tissue; o, organoids; GB, gallbladder Nomenclature for tissue samples: ‘t_’ is followed by patient ID. The patient ID includes a letter code indicating pathology group (n/x/p/m) followed by 2-digit patient serial number. External transcriptomic data was sourced from a published article^1^ for three pre-invasive malignant (adenoma) patients. SRR19139557, SRR19139556, SRR19139558 patient IDs in the published article have been indicated here as et_p01, et_p02 and et_p03, respectively. Nomenclature for organoid samples: ‘o’ is followed by a number to indicate different organoid line numbers developed from the same patient tissue but either using different types of growth media or different tissue dissociation methods for seeding cultures. Patient ID of the respective source tissue is indicated at the end. For detailed description of the samples, see Supplemental Table S6.

Gene expression of tissues and organoids were analysed using unsupervised hierarchical clustering based on top 1000 highly variable genes [Supplemental tables S4A, S4B, S5A and S5B]. For primary gallbladder tissues, samples from malignant pathology group (*m*) (adenocarcinoma, adenosquamous carcinoma, squamous cell carcinoma and small cell carcinoma) clustered separately from non-malignant (*n*) group (normal, chronic cholecystitis, cholesterolosis, acute on chronic cholecystitis). XGC (*x*), with one exception, clustered adjacent to the *m* group. We categorised intra-cholecystic papillary neoplasm (ICPN) as a pre-invasive neoplasm (*p*). We had only one sample (t_p08) of this very rare pathology. As there are no clearly defined criteria to distinguish between gallbladder adenoma and ICPN, 3 samples (et_p01, p02, p03) with gallbladder adenoma were included from previously published data^21^ to increase the sample size of *p* group. The *p* group, with one exception, clustered with the *m* and *x* cohorts [Figure 7B top panel]. Likewise, in unsupervised hierarchical clustering of organoid gene expression the *m*, *x* and *p* cohorts clustered separately from the *n* cohort [Figure 7B bottom panel].

Tissue samples contain an admixture of gallbladder cholangiocytes with other cells while PDOs are cholangiocytes derived from primary tissue, grown and expanded in Matrigel^TM^ or Geltrex^TM^ under controlled ex vivo conditions, Unsurprisingly, in t-distributed stochastic neighbour embedding (t-SNE) analyses of gene expression, tissues clustered separately from the PDOs [Data not shown].

t-SNE analysis of tissues shows 3 distinct groups, one (I) with only *n*, the second (II) with mostly *n*, 2 *m* and 1 *x* and the third (III) with mostly *m* along with *x*, *p* and 3 *n* samples [Figure 7C left panel]. t-SNE analysis of PDO gene expression shows two distinct clusters, the first with the *n* and 1 *x* and the second with *m*, *p* and *x.* Within these pathology-defined clusters in PDOs, *n* but no other samples segregated according to culture conditions [Figure 7C right panel]. *x* and *p* cluster along with *m*, distinct from *n* samples both for tissues and PDOs, more prominently for the latter.

DESeq2 was used to identify genes with low variance in expression between tissues and PDOs. For *n* group tissues/PDOs, 10% of genes had comparable expression, increasing to 31% for XGC and malignant tissue/organoids [Supplemental Figure S5A]. The lower number of similarly expressed genes in *n* group samples is possibly due to the heterogeneity of the samples and different culture conditions [Figure 7C right panel]. We have shown earlier that PDOs derived from malignant and XGC samples retain the cytopathological features of the primary tissue [Figure 5, Supplemental Figures S2, S3]. Thus, the higher proportion of genes commonly expressed between tissues and PDOs from *m* and *x* groups than *n* group, which comprise almost one-fifth of all genes, are likely to be associated with morphological phenotypes [Supplemental Figure S5A]. 2170, 1960 and 531 genes were exclusively detected in *m*, *x* and *n* groups, respectively. Hallmark pathway enrichment using these genes showed E2F targets, G2M checkpoint, mitotic spindle, Myc targets_v1 and mTORC1 signalling enriched in the *m* group. Oxidative phosphorylation, Myc tagets _v1 and unfolded protein response pathways were enriched in the *x* group [Supplemental Table S7 and Figure 7D]. In the 1995 genes not specific for any one pathology group, 1246 (62%) were shared exclusively between the *m* and *x* groups - higher than the exclusively shared gene expressions between *m* and *n* (10%) or *n* and *x* (18%) groups. This suggests the genes expressed by *x* pathology was closer to *m* than to *n* group pathologies [Supplemental Figure S5B. The histological characteristics, hierarchical clustering and commonality of gene expression of XGC organoids with GBCOs suggest that the intensive chronic inflammatory XGC occupies an intermediate state between normal and malignant GB.

Taken together, the data shows the developed PDO lines can be used as suitable models for studying gallbladder diseases.

## Discussion

For the first time, we report the creation of a characterised repository of primary gallbladder tissues and living GCOs/GBCOs developed from freshly processed and cryo-preserved primary cholangiocytes derived from a wide range of gallbladder pathologies, including invasive malignancies, pre/non-invasive neoplasms (ICPN, adenoma), inflammatory diseases such as chronic cholecystitis or the rare XGC, and pathologically normal gallbladders. While GCOs from normal gallbladders have been successfully generated ^22^, there has been limited success with other GB pathologies and in particular GBC.^8,10–12^ A rate limiting step is access to fresh tissue. Though GBC is prevalent in our region, most patients have inoperable disease and in a third the diagnosis is incidental with the patient presenting with a prior cholecystectomy.^5^ Even at a dedicated cancer centre like ours, patients with GBC are managed by several different departments. We developed an integrated team with a designated patient liaison coordinator. Over time a cost-effective and streamlined workflow to obtain samples was established. GCOs/GBCOs were generated from fresh or cryopreserved primary cholangiocytes derived from surgically resected gallbladders, but not from core biopsies. The latter were primarily from hepatic or omental metastases which were often necrotic and lacked cellularity ^23^. Often the amount of surgically resected tissue is insufficient for organoid generation. That along with presence of large portion of fibrous connective tissues are possible contributing factors leading to the reported success rates of 20% or lower to generate GBCOs.^8,9,24^ For small tissue samples, the use of optimised enzymatic digestion mix with collagenase/dispase and DNaseI (supplemented with a ROCK inhibitor Y-27632) to isolate cholangiocytes from primary tissue and passaging degenerating organoids using TrypLE digestion (supplemented with Y-27632) improved success rates of GCO/GBCO generation and propagation to 58% (69/119) overall and 52% (10/19) for the *m* group pathologies in the 2022-23 period [Figure 3E and Supplemental table S2]. We used two different culture conditions to generate gallbladder PDOs, one with GSK3 inhibitor CHIR99021 to stimulate^9^ and the other in which this was omitted and DKK-1 added to inhibit^15^ the canonical WNT pathway respectively. Both were equally effective in generating PDOs from different gallbladder pathologies [Figure 3C]. Both GCOs and GBCOs were amenable to long term propagation with retention of cytopathological features in the GBCOs.

Like any disease model, these GCO/GBCO models have their own share of limitations. Currently we do not have GBCOs developed from metastatic or advanced stage of GBC as these tumours are inoperable and GBCO could not be grown from core biopsies. Some discordance was observed in certain marker protein expressions e.g. CK20 and HER2 [Figure 6E, Supplemental Figure S4C]. Thus, GCOs/GBCOs do not necessarily reflect the entirety of spatially heterogenous pathological changes occurring in the primary tissue as previously reported for HER2 expression in breast cancer.^25^ We recommend verification of specific organoid lines grown in any laboratory for expression of relevant markers, functionality and histological recapitulation prior to further experimentation. While transcriptomic analyses confirmed the developed GCOs/GBCOs preserved SNPs from the source tissues, we could not estimate the proportion of deleterious mutations preserved in the GBCOs as we had too few (m52 and m53) samples from the malignant group with patient-matched mRNAseq data from primary tissues and derived organoids for reliable analysis. Considering the rarity of surgically resected malignant gallbladders, mutational profiling would benefit from a multicenter study.

In unsupervised hierarchical clustering analysis XGC samples grouped with invasive malignant and pre-invasive neoplasms rather than other non-malignant diseases. This pattern was more pronounced in PDOs than in tissues, suggesting that PDOs better represent the changes in cholangiocytes from various pathologies. Notably, higher proportion (62%) of gene expression commonality between *x* and *m* group suggests XGC’s potential to transition to pre-neoplastic disease state. XGC is thought to start as biliary obstruction with rupture of Rokitansky Aschoff sinuses leading to bile extravasation. The resulting xanthogranulomatous inflammatory reaction can extend into adjacent tissue, mimicking an invasive neoplasm. Chronic inflammation is the main process that drives the genesis of GBC. Currently, two major histogenic sequences are proposed for GBC pathogenesis. More commonly metaplasia-dysplasia-carcinoma and less commonly adenoma/ICPN-carcinoma sequence, both preceded by long term chronic inflammation^4^. Previously, based on IHC expression of p53, PCNA and beta-catenin it has bene speculated that XGC is unrelated to GBC ^26^. In our PDO models, sustained chronic inflammatory conditions such as XGC appear to lead to activation of certain hallmark pathways associated with oncogenesis, without the presence of considerable dysplasia. This supports the hypothesis that a chronic inflammatory process in the GB leads to histogenic progression. This GCO/GBCO organoid bank which contains PDO from normal, inflamed, XGC, ICPN and malignant GBs can facilitate investigations into the processes that give rise to GBC.

Given the high prevalence of this mostly fatal disease in our region but with the practicality of obtaining fresh tissue, organoid banks such as ours are a unique resource for those who investigate disease pathogenesis and wish to identify novel therapies. Further strengthening of the bank requires multi-institutional collaborations across geographical boundaries.

## Resource Availability

### Materials availability

No new reagents have been generated in this study. Banked organoid lines are available on request to lead contact against a completed material transfer agreement and reasonable compensation for processing and shipping.

### Data and code availability

No original code has been developed in the current study. mRNAseq data is available at NCBI SRA (PRJNA1209630).

## Acknowledgement

This work is part funded by a Wellcome-DBT India Alliance Margdarshi Fellowship (IA/M/12/500755) and a Tata Consultancy Services Foundation core grant to VS. Ministry of Education, India PhD Fellowship to AkD (Ankita Dutta). We thank Cancer Genomics Laboratory, Clinical Proteomics Unit, IT unit, Clinical Research Unit, Biobank (TiMBR), admin and house-keeping teams at TTCRC. We appreciate Pathology, Radiology, GI-HPB Operation Theatre teams and nurses at TMC for their support. We express gratitude to the patients and family for donating clinical samples.

## Author contribution

Conceptualisation: VS and AD (Anindita Dutta); Experiment design: VS, AD, DGS; Performing experiments and data collection: AkD, NC, SC, DB, SJS, DGS; Coordination, patient consenting and clinical sample collection: PB, AK, ASR, MKR, SB, SS; Data analysis and interpretation: AkD, NC, AVS, DGS, AD, US, RS, PR, VS; Supervision: VS, DGS, AD, RS, US, TD, DG, SK, MKR, SB, SS; Writing original draft: DGS, AkD, NC, AVS; Review and critical editing: DGS, PR, VS; Funding acquisition: VS;

## Methods

Detailed methods are provided in the online version of this article which include the following:

o Patient identification and sample collection
o Extractions of biomolecules and quality checks
o Seeding and maintenance of GCOs
o Cryopreservation and retrieval of primary cells and organoids
o Histological analysis and immunohistochemistry of organoids
o Immunofluorescence
o Functional assays
o qRT-PCR analysis
o mRNA sequencing and analysis
o Data collection and management

## Materials and methods

### 1. Patient identification and sample collection

Patients with gallbladder diseases (malignant or non-malignant) were identified from the hospital electronic medical records (EMR) at the time of their visit to out-patient departments (OPD) of TMC between September 2019 to December 2023. Informed consents were obtained from patients who were eligible to participate in the study. Ethical approval was obtained from Institutional Review Board (EC/TMC/65/16 and 2021/TMC/216/IRB3). Tissue samples were obtained from patients who underwent gallbladder removal surgery or core needle biopsy procedure for suspected GBC at TMC. Samples were collected in 250mL (surgical) or 10mL (core biopsy) cold sterile phosphate-buffered saline (PBS) supplemented with 5ng/mL EGF (AF-100-15, Peprotech), 2μM Y27632 dihydrochloride (1254, Tocris), 1X penicillin/streptomycin (15140-122, ThermoFisher Scientific), 5μg/mL amphotericin B (Amphotret) and 5μg/mL fluconazole (Cipla). Also see Supplemental table 1 for alternative compositions for tissue transport media. The specimens were washed 3 times with sterile 1X PBS before downstream processing. Approximately 30mg of tissue was either snap frozen in liquid nitrogen (LN2) and stored at -80°C. Alternatively, similar amount of tissues was stored in 1mL RNAlater (AM7021, ThermoFisher Scientific) at 4°C or in TRI reagent (T9424, Sigma-Aldrich) at -80°C for RNA extraction. For DNA extraction, 30mg of tissue was stored in 180μl ATL buffer (939011, QIAGEN) at room temperature. The rest of the tissues (surgically resected samples) were processed for developing GCOs.

### 2. Extractions of biomolecules and quality checks

#### 2.a. DNA

Genomic DNA was isolated from the surgical tissue, snap-frozen tissue, formalin-fixed paraffin-embedded (FFPE) tissue, and GCOs. Lysis buffer was prepared by adding 20µL of Proteinase K (19131, Qiagen) into 180µL of ATL buffer (939011, Qiagen). Approximately 30mg of snap frozen tissue was crushed in LN2 and added to the 200µL of lysis buffer. For extracting DNA from freshly collected gallbladder surgical tissue samples, tissue (∼30mg) was homogenised in 200µL of lysis buffer using the battery-operated motorised pellet pestles (Z359971, Sigma-Aldrich). For lysing GCOs, 200µL of lysis buffer was added to the harvested and cold PBS washed organoid pellet. Next, for all types of samples lysis was performed at 56°C in shaking condition (660rpm) for overnight (16-18 hours). Then genomic DNA was isolated using the QIAamp® DNA Mini Kit (15304, Qiagen). For genomic DNA extraction from FFPE tissues, eight units of 10µm sections were used for each preparation and extraction was performed using the QIAamp DNA FFPE Tissue Kit (56404, Qiagen) following the manufacturer’s protocol with some modifications. Briefly, paraffin was removed with 1mL xylene (534056, Merck) treatment for 1 hour at 660 x g at 56°C shaking condition followed by the centrifugation at 17,000 x g at room temperature for 2 minutes for twice. Next, pellet was incubated for 30 minutes in 500µL of the nuclease-free water at 95°C for 15 minutes. This step was repeated twice followed by the ethanol wash mentioned in the manufacturer’s protocol. Then overnight lysis was done in the lysis buffer. Rest of the protocol was followed as recommended by the manufacturer. Freshly collected gallbladder tissue can be stored in 180µL ATL buffer at room temperature for a maximum of 30 days before proceeding to lysis.

Extracted DNA quantity was assessed with the Qubit® 4.0 Fluorometer (Q33216, Life Technologies) by using the Qubit™ dsDNA HS (Q32851, ThermoFisher Scientific) and Qubit™ dsDNA BR Assay Kit (Q32850, ThermoFisher Scientific). Extracted DNA quality was checked by gel electrophoresis run using 1% agarose (16500500, ThermoFisher Scientific) gel in 1X TBE buffer (574795, Sigma-Aldrich).

#### 2.b. RNA

RNAlater-stabilised tissues (25-30mg for surgically resected tissues and 2-3mg for core biopsy samples) or freshly harvested GCOs (from 2 confluent wells of 24-well dish) were transferred to the 1mL RLT lysis buffer (79216, Qiagen) freshly supplemented with 10μl 2-Mercaptoethanol (M3148, Sigma-Aldrich). Total RNA was extracted from the surgically resected gallbladder tissues (mucosal layer) using RNeasy mini kit (74104, Qiagen). For RNA extraction from core biopsy tissues and GCOs, RNeasy micro kit (74004, Qiagen) was used. Briefly the tissues or organoids were lysed by homogenizing with a battery-operated motorised pellet pestles for a maximum of 10min. The lysates were centrifuged at 10,000 x g for 3 minutes and the supernatants were processed for RNA extraction following the respective kit manuals including the on-column DNase digestion. RNA extraction was alternatively performed using TRI reagent. For this method, the samples (tissues or GCOs) were stored in 1mL TRI reagent at -80°C until extraction was performed. Immediately prior to the extraction, the frozen samples were brought to room temperature and lysed by repeated pipetting in TRI reagent. After lysis, RNA isolation was carried out following manufacturer’s protocol. DNase digestion was not performed for RNA samples isolated using TRI reagent. All the RNA samples were stored at -80°C in small aliquots. Tissue can be stored in RNAlater^TM^ at 4°C for a maximum of 30 days before proceeding to lysis.

One of the aliquots from each RNA sample was used for quantification by Qubit® 4.0 Fluorometer and RNA quality was analysed by automated gel electrophoresis analysis (4200 TapeStation System, Agilent). Qubit™ RNA Broad Range Assay kit (Q10211, ThermoFisher Scientific) was used for Qubit measurement. High Sensitivity RNA Screen Tape and corresponding sample buffer and ladder system (5067-5579, 5067-5580, 5067-5581, Agilent) were used for TapeStation analysis.

#### 2.c. Protein

Protein was extracted from the either snap frozen tissue or fresh tissue, and GCOs. Snap frozen tissue (∼30mg) was crushed in LN2 and 100μl of lysis buffer (A40006, EasyPep™ Mini MS Sample Prep Kit, ThermoFisher Scientific) was added to it. For freshly collected surgical tissue, gallbladder mucosa sample (∼30mg) was isolated and 100μl of lysis buffer was added for protein extraction. Tissue was homogenised with the battery-operated motorised pellet pestles. For extracting proteins from GCOs, organoids from four 80-90% confluent wells of a 24-well dish were snap frozen in LN2 and stored at -80°C. EasyPep™ Mini MS Sample Prep Kit was used for the protein extraction from the above-mentioned samples. Protein was quantified using Pierce™ BCA Protein Assay Kit (23227, ThermoFisher Scientific). Protein integrity was checked by SDS-PAGE electrophoresis.

20µg of total protein was used for the sample preparation for mass spectrometric analysis using EasyPep™ Mini MS Sample Prep Kit. The peptides were reconstituted in 100 µL of 0.1% formic acid (94318, Honeywell) in MS-grade water (900682, Sigma-Aldrich), loaded onto NanoLC coloumn (Eksigent Technologies) for liquid chromatography and mass spectrometric analysis was performed using Sciex TripleTOF 6600+ system.

### 3. Seeding and maintenance of GCOs

Cells were dissociated from the tissue either by mechanical scraping, collected in cold William’s E media (A1217601, ThermoFisher Scientific); or by enzymatic digestion of tissue using 10μg/mL collagenase-dispase (11097113001, Roche), 3.2μg/mL DNase-I (D5025, Sigma-Aldrich), 1X penicillin/streptomycin, 10μM Y27632 dihydrochloride in 10mL advanced Dulbecco’s modified eagle medium/F12 (ADMEM/F12) (12634010, ThermoFisher Scientific) for 60–90 minutes (depending on tissue amount) at 37°C at 71rpm in a shaking incubator (Incu-Shaker Mini, Benchmark). The dissociated cells were filtered through a 70μm cell strainer (352350, Corning) and centrifuged at 450 x g for 5 minutes followed by removal of red blood cells contamination using RBC lysis buffer (11814389001, Roche) for 5 minutes at 4°C. The pellet after centrifugation was washed thoroughly with cold media. Cells were resuspended in ice-cold 20μL growth media and 40μL matrix - Matrigel^TM^ (356237, Corning) or Geltrex^TM^ (A1413202, ThermoFisher Scientific). 20,000 – 50,000 cells per well were seeded in a 24-well dish (3526, Corning). Matrix was allowed to be solidified at 37°C for 5-7 minutes. The plate was inverted upside down and kept for additional 15 minute for the cells to be entrapped within the dome of matrix. Each well was then covered with 1 mL of organoid growth media either in type A or type B organoid growth media [Supplemental table S3 for composition] and kept in 37°C incubator with 5% CO_2_ and 95% relative humidity (Galaxy 170S, Eppendorf).

Once GCOs were formed, the culture was maintained in respective growth media. The medium was replaced with fresh growth media every 3-4 days. GCO cultures were passaged every 7 to 10 days based on their growth. For passaging, GCOs were harvested in cold cell recovery solution (354253, Corning) and incubated for 30 minutes on ice to remove matrix. Harvested GCOs were mechanically broken into smaller pieces by repeated pipetting. For GCO cultures not reaching 70-80% confluence within 7 to 10 days or showing degeneration, harvested organoids were further dissociated using 200μl TrypLE Express^TM^ (12605-010, ThermoFisher Scientific), supplemented with 10μM Y27632 dihydrochloride and incubated at 37°C for 5 minutes at 71rpm in a shaking incubator. The digestion was stopped with cold ADMEM/F12 media. GCOs were next re-seeded in fresh matrix and overlaid with media for maintaining the culture. Images of the growing GCOs were captured under Nikon Eclipse TS2 microscope at regular interval.

### 4. Cryopreservation and retrieval of primary cells and organoids

Cell counting was performed using haemocytometer cell counting chamber upon trypan blue staining. A minimum cell viability of 40% was preferred for cryopreservation. 0.8 to 1×10^6^ viable primary cells were resuspended in 1mL freezing media (FBS: DMSO – 9:1) supplemented with 10µM Y27632 dihydrochloride. The vials were transferred to -80°C in gradient cell freezing container (5100-0001, ThermoFisher Scientific) and then to the LN2 storage within 2 days.

Whole GCOs (preferably <200µm average diameter) from two 70-80% confluent wells of a 24-well dish were harvested in 1mL cold Cell Recovery Solution, incubated on ice and washed with cold PBS to remove traces of matrix. The GCOs were pelleted at 100 x g for 5 minutes. Alternatively, organoids (>200µm average diameter) were enzymatically dissociated to small cell clusters using TrypLE Express as discussed for passaging. A minimum cell viability of 60% was preferred for cryopreservation. 0.8 to 1 ×10^6^ viable cells were cryo-preserved per vial. TrypLE-digested GCOs or intact GCOs were then resuspended in 1mL Recovery Cell Culture Freezing Medium™ (12648010, ThermoFisher Scientific) supplemented with 10µM Y27632 dihydrochloride and stored as discussed for cryopreservation of primary cells.

For generating organoids from banked samples, the cryo-preserved primary cells or organoids were retrieved from LN_2_ and immediately thawed at 37°C water bath (HH-2J, NeuProCell) with mild stirring until only a small part remained frozen. The entire suspension was then transferred into 10mL pre-warmed ADMEM/F12 or William’s E media supplemented with 10μM Y27632 dihydrochloride. The primary cells or enzymatically digested organoids were centrifuged at 450 x g and whole organoids at 100 x g for 5 minutes at room temperature. The pellet was resuspended in matrix and seeded into matrix in the 24-well culture dish as discussed previously. Prior to seeding, the primary cells were treated with 3.2μg/mL DNase I (in 2mL of warm ADMEM/F12 media supplemented with 10µM Y27632 dihydrochloride) for 15 minutes at 37°C at 71rpm in a shaking incubator. The tips of the sterile 1000µL micropipette tips were cut before use to increase the barrel diameter to prevent the organoids from breaking or shearing during the entire procedure for cryo-preservation and retrieval of whole organoids.

### 5. Histological analysis and immunohistochemistry of organoids

GCOs were harvested in cold cell recovery media using cut tips to avoid breaking them. Next, they were fixed in 10% Neutral Buffered Formalin (NBF) at 37°C for 15 minutes and washed with cold PBS for two times. For each wash, 5-10 minutes of time was allowed for the GCOs to settle down on gravitational force only, centrifugation was avoided. Washed GCOs were mixed with 50μL of fresh plasma and equal volume of thromboplastin (HemosIL RecombiPlasTin 2G, 0020002950, Instrumentation Laboratory Co.) to form a coagulum. Then each coagulum was transferred to a piece of biopsy wrap paper (3801090, Leica Biosystems) and stained with one drop of 1% (w/v, in 100% ethanol) eosin Y solution (230251, Sigma-Aldrich) followed by wrapping inside the biopsy wrap paper. Each of the folded biopsy papers containing the eosin stained organoid coagulum or primary tissue pieces was put inside a biopsy cassette (IP-BIOPSY-CASSETTE-III, Leica Biosystems) and incubated in 10% NBF for 4 hours at room temperature. The FFPE blocks were made using Leica Biosystems Heated Paraffin Embedding Station (Histocore Arcadia H). 5µm or 3µm sections from the FFPE blocks were cut using microtome (Histocore Biocut, Leica Biosystems), flattened on warm (45°C) water, collected onto positively charged glass slides (BOND-PLUS-SLIDES, Leica Biosystems) and dried for haematoxylin-eosin (HE) staining and immunohistochemical (IHC) analysis, respectively. For IHC, antibodies were used against CK7 (1-CY163-07, Diagomics, 1:100), CK20 (MU315-UC, BioGenex, 1:50), and HER2 (790-2991, Roche) markers. Antigen retrieval and staining was performed in automated system using Leica Bond-Max (49.0051, Leica Biosystems) for CK7 and CK20 and Ventana benchmark ultra (Ventana) for HER2. IHC scoring for HER2 marker was performed following the guidelines for gastroesophageal adenocarcinoma by ASCO/CAP 2016. For CK7 and CK20 markers, a cumulative score was determined by multiplying an intensity sore and a distribution score, both individually ranging from 1 to 3 or 0 for no expression. Score 3: Strong expression intensity or diffuse spread, score 2: moderate intensity or patchy expression, score 1: weak intensity or only focal expression, score 0: no expression. Thus, cumulative score ranged between 0 and 9. For plotting in the heatmap, cumulative score of 0-3 was considered as no or low expression, 4-6 as intermediate and 7-9 as high expression. Images of the stained sections were captured by Leica DMi8 in bright-field mode.

### 6. Immunofluorescence

GCOs embedded in matrix were fixed with 4% paraformaldehyde (28908, ThermoFisher Scientific) for 20 minutes at 4°C and the auto-fluorescence was quenched using 50mM ammonium chloride (A9434, Sigma-Aldrich). Organoids were permeabilised and non-specific binding of antibodies was blocked using 10% goat serum (G9023, Sigma-Aldrich) and 0.1% Triton X-100 (X100, Sigma-Aldrich) in PBS for 1 hour at room temperature. Organoids were then incubated with primary antibody solutions; CK19 (ab52625, Abcam) and Mucin-5B (HPA008246, Sigma-Aldrich) - dilution 1:100 each, overnight at 4°C. Post washing suitable secondary antibody (A21428, Invitrogen) was used overnight at 4°C (dilution – 1:1000). GCOs were counterstained with Hoechst 33342 (62249, ThermoFisher Scientific) for 10 minutes at room temperature. Images were acquired using a confocal microscope A1RHD (Nikon).

### 7. Functional assays

GCOs were harvested and seeded in a 10µL matrix dome in a 96-well plate (167008, ThermoFisher Scientific). All the functional assays were performed after 5-7 days of seeding.

#### 7.a. Epithelial barrier assay

Epithelial barrier assay was performed as previously described by *Sepe et al*. GCOs were incubated with PBS containing 1mg/mL fluorescein isothiocyanate dextran (FITC-dextran, 46944-F, Sigma-Aldrich) and incubated at 37°C. Images were captured at 470nm wavelength on Nikon Eclipse TS2 after 30 minutes of incubation. 2mM ethylene glycol-bis (β-aminoethyl ether)-N, N, N′, N′-tetraacetic acid (EGTA, E3889, Sigma-Aldrich) was added and incubated at 37°C. Images were captured after 30 minutes.

#### 7.b. Pump activity assay

GCOs were incubated with fresh medium containing 100μM Rhodamine 123 (R8004, Sigma-Aldrich) for 5 minutes at 37°C and washed with 1X PBS. GCOs were harvested using cold Cell Recovery solution using 1000μl cut tips (to keep the whole organoids intact) and washed to remove traces of matrix. Images were captured at 470nm wavelength on Nikon Eclipse TS2.

To show that the rhodamine 123 uptake was due to the ABCB1 pump, fresh medium containing 10μM verapamil hydrochloride (V4629, Sigma-Aldrich); MDR (multi-drug resistant) inhibitor was added and incubated for 30 minutes at 37°C and the assay repeated.

#### 7.c. Alkaline phosphatase assay

The substrates to detect alkaline phosphatase were prepared according to the manufacturer’s instructions BCIP/NBT Color Development Substrate (5-bromo-4-chloro-3-indolyl phosphate/ nitro blue tetrazolium) (S3771, Promega). GCOs were incubated with fresh colourless medium containing the substrates for 1 hour at 37°C. Images were captured using Leica DMi8.

#### 7.d. Gamma glutamyl transferase activity

Post removal of matrix GCOs were enzymatically digested using TrypLE Express as discussed for passaging. 2000 cells per well were seeded in a 10µL Matrigel dome in a 96-well plate. After 5-7 days of seeding GGT activity was measured in triplicates using the Gamma Glutamyl Transferase Assay Kit (colorimetric) (ab241029, Abcam). Absorbance readings were measured at 418nm using SpectraMax® M Series Multi-Mode Microplate Readers. The GGT activity was normalised against the total cell count post assay.

### 8. qRT-PCR analysis

cDNA was synthesised from 250ng of the total RNA using the High Capacity cDNA Reverse Transcription kit (4368814, ThermoFisher Scientific). The PCR product was quantified using NanoDrop 2000 spectrophotometer (ThermoFisher Scientific) and 10ng cDNA was used as a template for Real-time PCR. qRT-PCR was performed in triplicates using PowerUp™ SYBR™ Green Master Mix (A25742, ThermoFisher Scientific) on QuantStudio 7. All the primers used in this study were designed using Integrated DNA technologies (IDT) and used in the concentration of 10μM each. All the primers have a comparable efficiency of 90-105%. Differential gene expression relative to the reference gene (GAPDH) was calculated.

### 9. mRNA sequencing and analysis

For mRNA-seq library preparation, RNA samples with RNA integrity number equivalent (RIN^e^) value of >7.0 was considered. 200ng of total RNA was used from a fresh aliquot to prepare each cDNA library using TruSeq Stranded mRNA library prep kit (20020595, Illumina) following the kit manual. Briefly, poly-A containing mRNA molecules were purified using oligo-dT magnetic beads and they were fragmented into small pieces using divalent cations under elevated temperature. Using random primers and SuperScript II reverse transcriptase (18064022, ThermoFisher Scientific) the mRNA fragments were reverse-transcribed to first strand cDNA. Actinomycin D was added to prevent unwanted DNA-dependent synthesis. This was followed by second strand synthesis using DNA polymerase I and RNase H. Next, a single base 3’-adenylation to the blunt cDNA fragments was done to prevent self-ligation during adapter ligation step. Then single index or dual index adapters were ligated to the DNA fragments. Finally, the adapter ligated cDNA fragment library preps were enriched and stored at -20°C until further use. The cDNA library preps were quantitated by Qubit 4.0 Fluorometer using dsDNA high sensitivity Assay kit (Q32851, ThermoFisher Scientific). The library prep quality was tested by automated electrophoresis analysis (4200 TapeStation System, Agilent) using DNA 1000 high sensitivity assay kit (5067-1504, Agilent). 5µL of 4nM cDNA library from each prep was used for library denaturation. Sequencing run was performed in Illumina NextSeq 550 system (paired-end, 150 cycles x 2) upon preparing the sequencing sample using NextSeq 550Dx High Output reagent kit v2.5 (300 cycles) following the manufacturer’s protocol.

For data analysis, raw reads were first quality checked using FastQC. Then sequence alignment was performed using STAR alignment (v.2.7.8) and counts were read using human genome (GRCh38 Ensembl 90) as the reference. QC analysis of the aligned reads were performed using Samtools flagstat, CollectInsertSizeMetrics and CollectRnaSeqMetrics (Picard). Variants were called from the two-pass aligned RNA-Seq bam with the haplotyper caller from GATK. SNVs (Single Nucleotide Variants) from tissue and organoids were compared. Overlapping intervals in each tissue and matching organoid with a depth of at least 10 reads mapped with a quality of >= 20 were first identified. SNVs occurring in these intervals were annotated as PASS and with a genotype quality >= 30 in the tissues were then compared with the respective organoids to calculate the percentage of overlap in the pair.

Next the data were subjected to either exploratory analysis or differential analysis. For exploratory analysis, gene count data was filtered and only genes with at least 10 read counts in more than 25% of the samples were kept. The counts were transformed with variance stabilisation transform (vst) from the DESeq2 (v1.40.2) R package and the 1000 highly variable genes were scaled [Supplemental table S4A, S4B, S5A and S5B]. For scaling following formula was used. Scaled vst for each gene = (vst value – mean vst)/ standard deviation of vst. t-distributed stochastic neighbour embedding (t-SNE) analysis was done and hierarchical clusters were plotted with Pearson distance metric and Complete clustering method using hclust function.

Genes with unaltered expression (low variance) between tissues and GCOs specific to each pathology were identified using DESeq2 method with genes having at least 10 counts in 75% of the samples. Genes with expression |logFC| < 1 and adjusted.p-value > 0.5, between tissues and GCOs for a given pathology were considered as unaltered gene expression. Genes present in more than one pathology were removed to identify the exclusive gene expressions for each pathology. These genes were enriched for the Hallmark geneset from MsigDB v2022 dataset using ClusterProfiler (v4.8.3) and genesets with adjusted.p-value < 0.05 were considered to be significantly enriched for each pathology.

### 10. Data collection and management

Pathological diagnosis of the GB tissues was obtained from hospital electronic medical record. Pre-analytical variables linked to collected samples, such as, time of collection and time delay to receive sample at the biobank, were collected and recorded in LIMS software (Labvantage). Details of sample processing, including derivatives obtained from each sample and their storage at the biobank were also recorded in LIMS. Other details, like transport condition, visible appearance of tissue or any deviations from the standard operating procedure is recorded on hardcopy datasheets.

