## Supplemental Figures for "Insights into gallbladder cancer pathogenesis from a living organoid gallbladder cholangiocyte biorepository"

**Supplemental Figure S1**

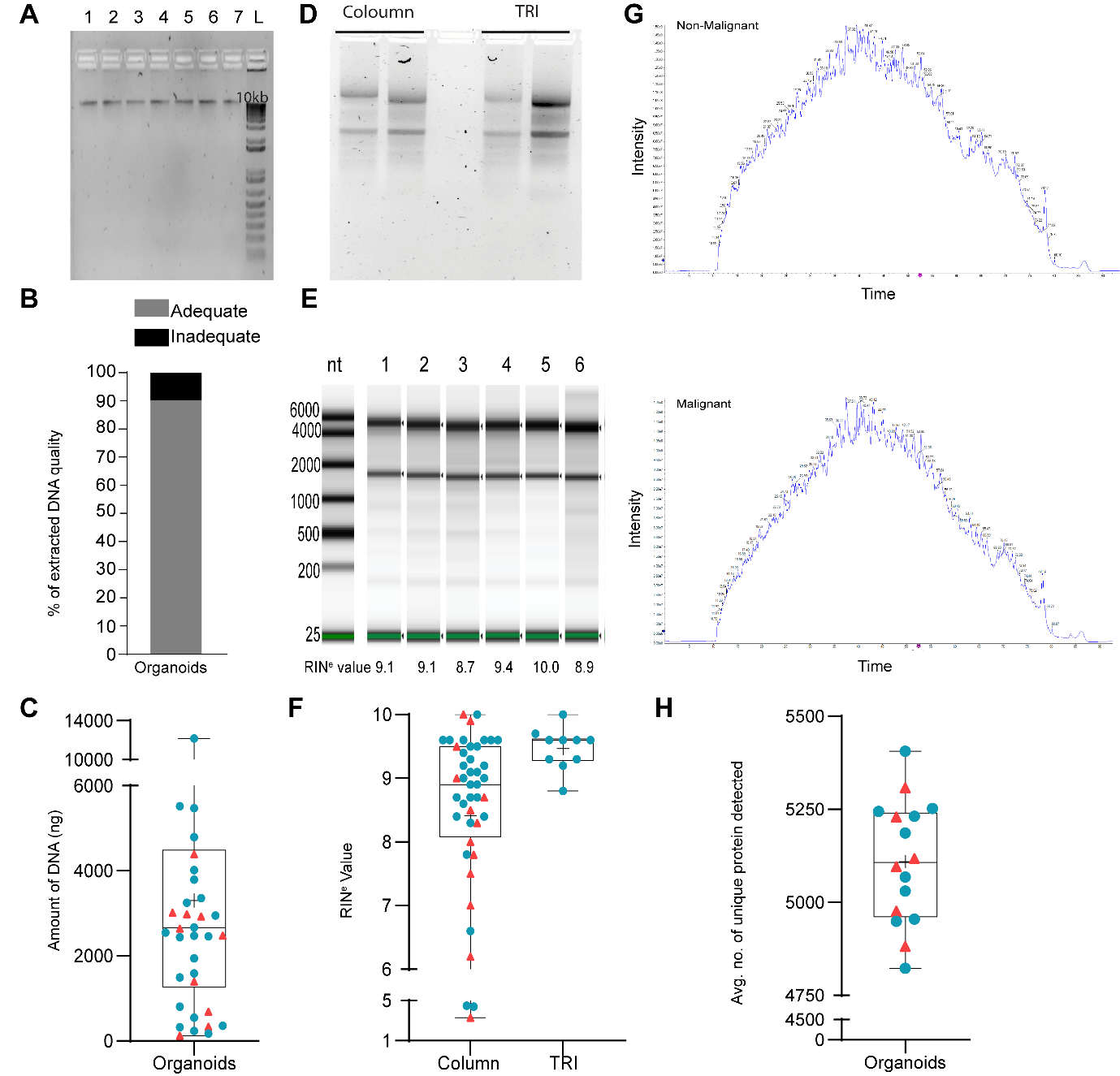

**Figure S1.** **Quality, purity and quantity of the extracted biomolecules from organoids were adequate for downstream high throughput experiments.**

**(A)** Representative image of 1% agarose gel electrophoresis analysis showing the integrity of gDNA samples extracted from freshly harvested organoids. The gDNA samples were extracted from GCOs and GBCOs derived from tissues with (1) normal, (2) chronic cholecystitis, (3) xanthogranulomatous cholecystitis with cholesterolosis, (4) adenoma, (5) ICPN, (6) adenocarcinoma and (7) adenocarcinoma in the background of ICPN, respectively. (**B)** Bar graph depicting proportions of organoid-derived gDNA sample integrity observed in agarose gel electrophoresis.  **(C)** Box and whisker plot depicting the distribution of total quantity of gDNA extracted from the organoid samples. **(D)** Representative image of RNA quality analysis by 1% agarose gel electrophoresis. **(E)** Representative images of column (RNeasy kit, Qiagen) purified organoid RNA samples run in automated gel electrophoresis for quality analysis. Organoids were derived from (1) normal gallbladder, (2, 3) acute on chronic cholecystitis, (4) XGC and (5, 6) chronic follicular cholecystitis with cholesterolosis gallbladder pathologies. **(F)** Box and whisker plots depicting the distribution of the RIN^e^ values of organoid RNA samples determined in RNA quality analyses by 4200 TapeStation System (Agilent). **(G)** Representative images of intensity-time chromatogram plots from LC-MS/MS analyses (DDA mode) of proteins extracted from organoids derived from XGC (top) and AdSqCa tissues (bottom). **(H)** Box and whisker plot depicting the distribution of number of unique proteins detected from each organoid-derived protein sample.

Box and whisker plots are as described in Fig 2. For all the agarose gel electrophoresis analyses, equal volume (2µl) of eluted gDNA or RNA were run in each lane.

TRI, TRI reagent^TM^; RIN^e^, RNA integrity number equivalent; nt, nucleotides; GCO, gallbladder cholangiocyte organoids; GBCO, gallbladder carcinoma organoids

**Supplemental Figure S2**

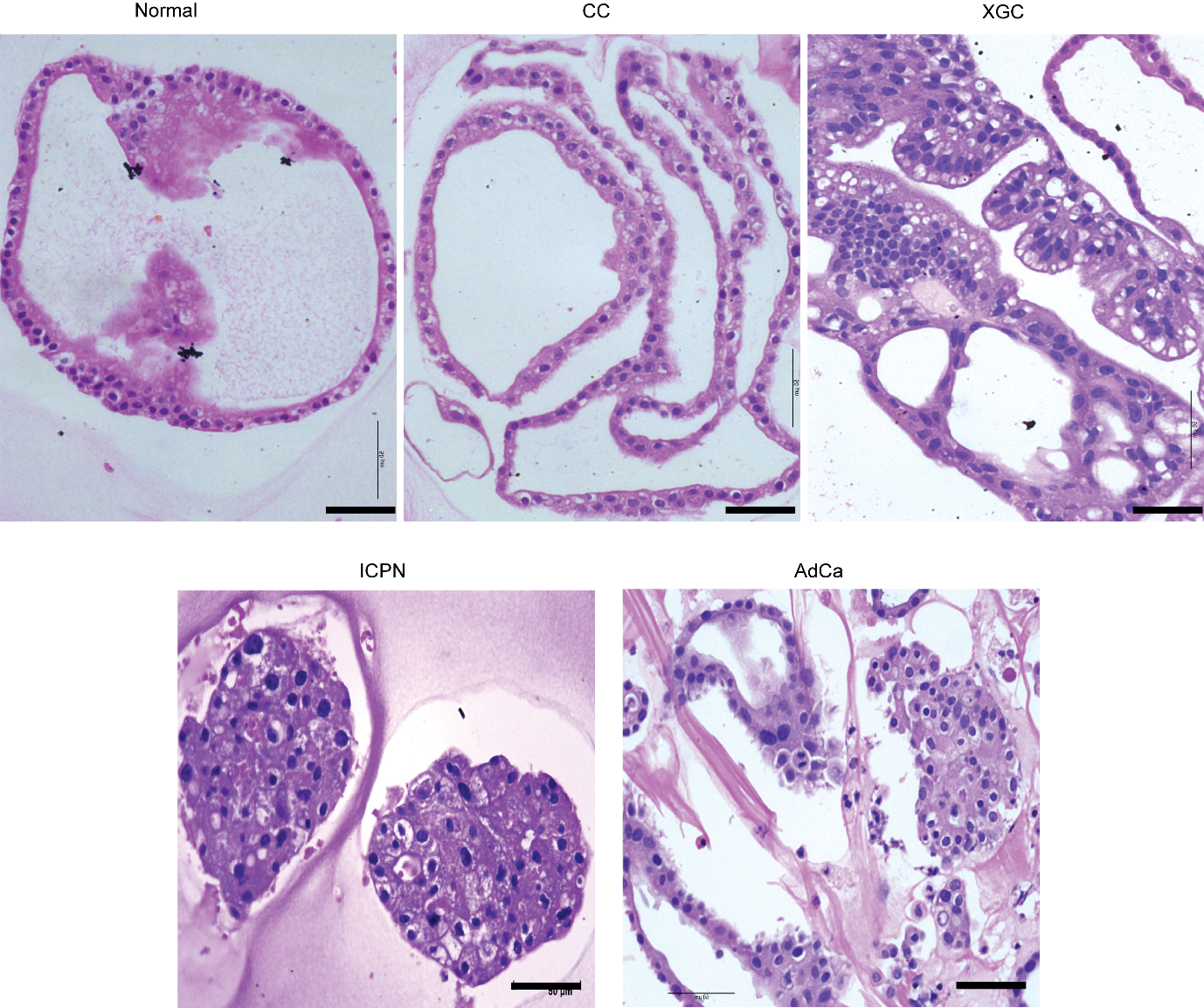

**Supplemental Figure S2: Higher magnification images of organoids showing preservation of pathology-specific architecture and cytology**

Higher magnification images of the areas marked by black rectangles in Figure 5B bottom row. Scale bars: 50µm. CC, chronic cholecystitis; XGC, xanthogranulomatous cholecystitis; ICPN, intracholecystic papillary-tubular neoplasm; AdCa, adenocarcinoma; HE, hematoxylin-eosin. Microscope: Leica DMi8 40X air objective, bright field mode.

**Supplemental Figure S3**

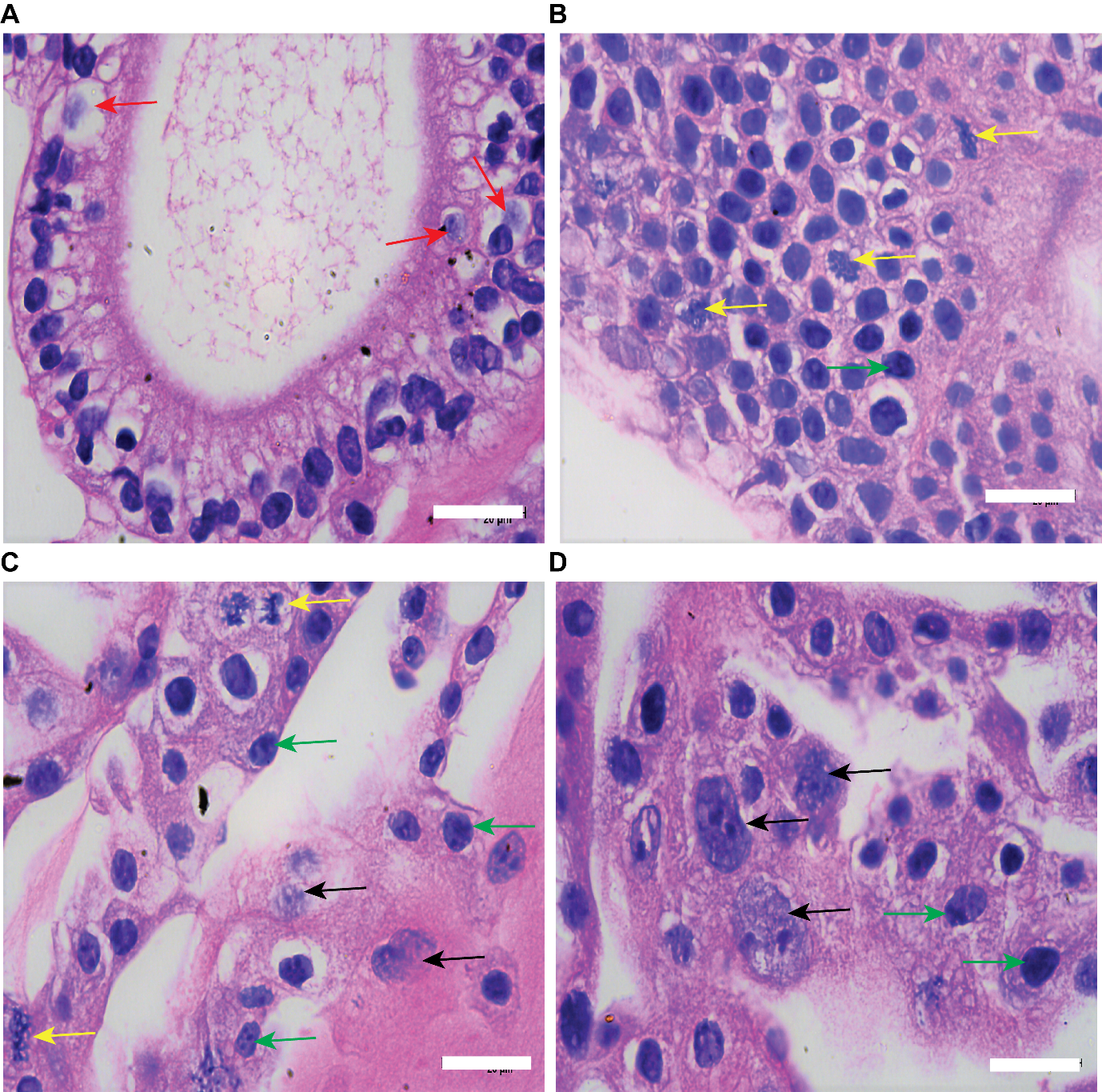

**Supplemental Figure S3: HE-stained images showing various cytological features observed in the developed organoids at higher magnification**

**(A)** Columnar epithelia including mucin secreting cells (red arrow) in GCO derived from normal gallbladder**. (B-D)** Higher magnification images of the HE-stained GBCOs developed from adenocarcinoma tissues exhibiting malignant cytological features like nuclear atypia, hyperchromasia, nuclear disarray, clumped chromatin (black arrow), mitotic cells (yellow arrow), multiple prominent nucleoli (green arrow). Scale bar - 20μm. GCOs, gallbladder cholangiocyte organoids; GBCO, gallbladder carcinoma organoids; HE, hematoxylin-eosin. Microscope: Leica DMi8, 100X oil objective, bright field mode.

**Supplemental Figure S4**

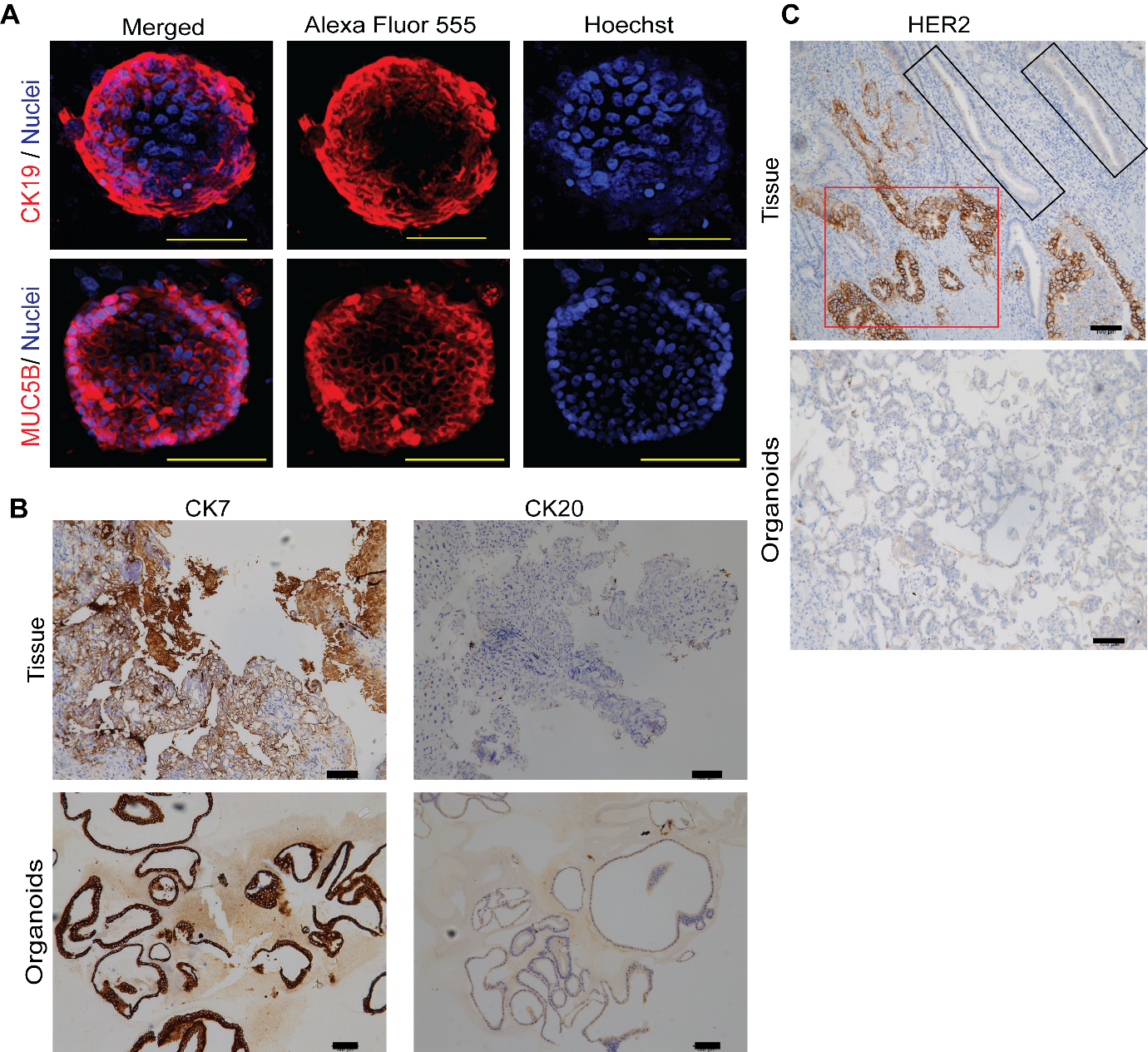

**Supplemental Figure S4: Organoid lines preserved the expressions of tissue protein markers except HER2**

Representative images of **(A)** immunofluorescence-stained GCOs showing expression of the marker proteins KRT19 and Mucin-5B. **(B)** Immunohistochemistry of matched tissues and organoid lines showing expression of CK7, CK20 in XGC and **(C)** HER2 in AdCa sample. Tissue sites with strong or no expression of HER2 marker have been indicated by red and black rectangles, respectively. Scale bars: 100µm (black), 50µm (yellow). Microscope: Nikon A1RHD (A), Leica DMi8 (B, C).

GCOs, gallbladder cholangiocyte organoids; XGC, xanthogranulomatous cholecystitis; AdCa, adenocarcinoma

**Supplemental Figure S5**

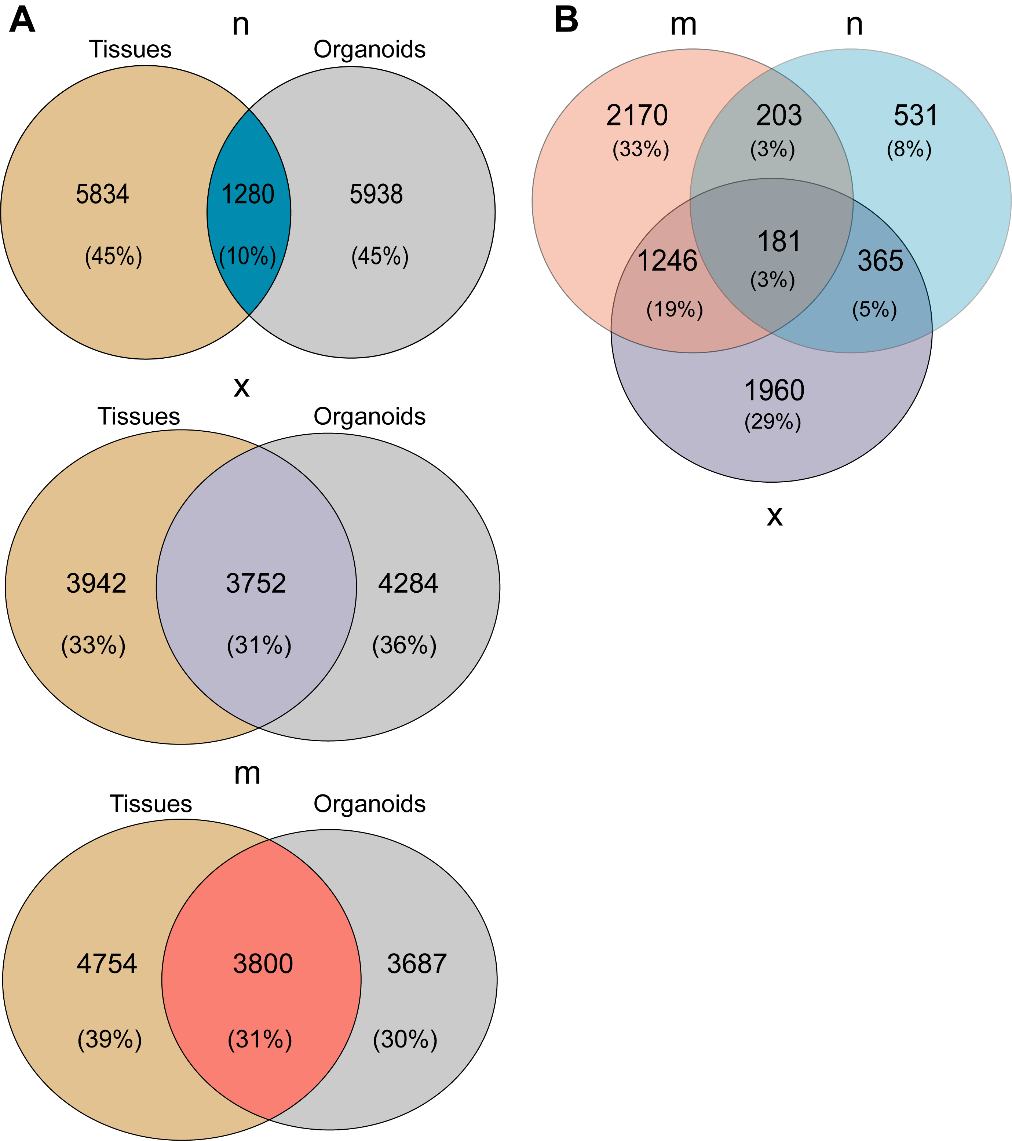

**Supplemental Figure S5: Organoid models developed from *m* and *x* group pathologies preserved higher proportion of tissue gene expression compared to *n* group**

**(A)** Venn diagrams representing the number of genes for which expressions were detected in tissues and organoids for each pathology group. Overlapped sections indicate the number of genes with unaltered expression (|log fold change|<1 and p>0.5) between tissues and organoids obtained by DESeq2 analysis across pathology groups- n (top panel), x (middle panel), m (bottom panel). **(B)** Venn diagram representing the number of genes with unaltered expression (|log Fold Change| < 1 and adj.p-value > 0.5) between tissues and organoids across pathology obtained by DESeq2 analysis.

All abbreviations and sample nomenclature used here are described in Figure 7.
